# Perineuronal net homeostasis by the NRG1–ErbB4 signaling for cognitive function

**DOI:** 10.64898/2026.11.20.768068

**Authors:** Bin Luo, Wenjun Zou, Jiawen Hou, Zhouyang Shen, Xinran Feng, Yiming Zhou, Huili Zhu, Lingli He, Wen-cheng Xiong, Lin Mei

## Abstract

Perineuronal nets (PNN) is the specialized extracellular matrix that enwraps the soma and proximal dendrites of neurons in particular interneurons. During neural development, PNN expression marks the closure of the critical period of visual cortical development. The roles of PNN in the adult brain and how its expression is regulated remain less understood. Here we show that enriched environment (EE) increases PNN in the prefrontal cortex (PFC) in adult mice with improved cognitive functions. Analysis of snRNA-seq data of EE and control PFC identified the neuregulin-1 (NRG1)–ErbB4 signaling as the pathway to regulate PNN expression. Ablation of ErbB4 or acute chemical genetic inhibition of its kinase activity reduced PNN and impaired cognitive function and prevented the promotion effect of EE. The expression of PNN and the core component aggrecan (ACAN) was increased in cultured interneurons stimulated with NRG1. ACAN mutation abolished PNN, weakened excitatory inputs and intrinsic excitability of ErbB4+ interneurons, and impaired cognitive functions. ACAN could undergo liquid-liquid phase separation to form condensates alone or together with other proteoglycans of the PNN. Finally, we demonstrate that restoring ACAN could ameliorate PNN and cognitive deficits in ErbB4 mutant mice. These results suggest that EE may improve cognitive functions by enhancing NRG1-ErbB4 signaling to increase PNN levels, identifying a signaling pathway critical to PNN homeostasis in adult animals

**Highlights:**

- Prefrontal PNN is required for enriched environment-increased cognitive function
- NRG1 promotes PNN expression by activating ErbB4 in interneurons
- PNN homeostasis requires ACAN which may act by phase separation
- ACAN in ErbB4⁺ interneurons enables EE to enhance PNN and cognitive function

## Introduction

The interstitial space in the brain is filled with the extracellular matrix that not only provides mechanical support for neurons but also is critical to their development and functions. One such structure is the perineuronal net (PNN) that enwraps the soma and proximal dendrites of interneurons and other cell types. PNN was first described by Camillo Golgi in 1898 as evidence to support the reticular theory but was viewed as a fixation artifact by Ramon y Cajal and had been ignored for many years (Celio et al., 1998). The PNN consists of a backbone of long hyaluronan chains, chondroitin sulfate proteoglycans (CSPG) such as aggrecan (ACAN), brevican (BCAN), neurocan (NCAN); CSPGs bind the hyaluronan, and link proteins (such as tenascin-R) to form a ternary complex. The functions of the PNN surrounding parvalbumin (PV)+ interneurons have been studied intensively. Its accumulation around PV+ interneurons marks the closure of the critical period of visual cortical development; enzymatic disruption of the PNN in visual cortex and barrel cortex unlocks juvenile plasticity and extends the critical period (Beurdeley et al., 2012; McRae et al., 2007; Pizzorusso et al., 2002). In addition, PNN levels continue to increase during development and plateau at P50 in the V1 area of the mouse visual cortex, for example (Slaker et al., 2016; Ye and Miao, 2013). These observations suggest that PNN may regulate brain functions in adult animals. Indeed, PNN disruption in the medial prefrontal cortex by chABC impairs cocaine-induced conditioned place preference memory and fear conditioning (Banerjee et al., 2017; Gogolla et al., 2009; Hylin et al., 2013; Slaker et al., 2015). On the other hand, chABC treatment was shown to enhance cognitive flexibility in reversal learning (Happel et al., 2014) memory performance (Cope et al., 2022; Domínguez et al., 2019; Romberg et al., 2013; Xue et al., 2014), recognition memory (Romberg et al., 2013) and spatial working memory (Anderson et al., 2020). The pattern of PNN of PV+ interneurons may encode long-term memory (Lev-Ram et al., 2024; Tsien, 2013) PNN can also protect neurons from oxidative stress and excitotoxicity by buffering local ion concentrations (Cabungcal et al., 2013; Härtig et al., 1999; Morawski et al., 2004). In accord, PNN appeared to be disrupted in regions affected by Alzheimer’s disease (Baig et al., 2005). However, roles of PNN in the adult brain are less understood.

PNN formation during development is a dynamic process. Rearing rats without visual stimuli in the dark from birth prevented the formation of PNN around PV+ interneurons; on the other hand, reintroducing the animals to a normal light/dark environment could restore PNN (Pizzorusso et al., 2002). In accord, reducing PV+ interneuron’s activity leads to disassembly of its PNN (Devienne et al., 2021). These results suggest that PNN formation is regulated by neuronal activity of PV+ interneurons. Moreover, PNN maturation requires homeoprotein Otx2 whose accumulation is enhanced by visual experience (Sugiyama et al., 2008). In addition, Sema3A accumulates in the PNN also in an experience-dependent manner and its presence in the mature visual cortex inhibits plasticity (Boggio et al., 2019). The formation and remodeling of the PNN requires the enzymes MMP (matrix metalloproteinase) and ADAMTS (a disintegrin and metalloproteinase with thrombospondin motifs), both being able to degrade CSPGs like ACAN and BCAN (Nakamura et al., 2000; Stanton et al., 2005; Tortorella et al., 2000). However, it remains unclear how the PNN is regulated in the adult brain, a glaring gap in our understanding its functions.

Neuregulin 1 (NRG1) is a trophic factor that acts by activating the receptor tyrosine kinase ErbB4 in the brain. In the cortex, NRG1 is produced mainly by pyramidal neurons in an activity-dependent manner whereas ErbB4 is specifically present in interneurons (Liu et al., 2011; Mei and Nave, 2014; Yao et al., 2023). The NRG1-ErbB4 signaling is critical to GABAergic circuit assembly by promoting interneuron migration, differentiation and synapse formation (Flames et al., 2004; Krivosheya et al., 2008; Ting et al., 2011; Yau et al., 2003; Yin et al., 2013b). PV+ interneuron-specific ablation or chemical genetic inhibition of ErbB4 disrupts GABA release and thus synaptic plasticity (Tan et al., 2018; Woo et al., 2007) and impairs various brain functions including working memory, attention, social novelty and fear conditioning (Bi et al., 2015; Del Pino et al., 2013; Lu et al., 2014; Robinson et al., 2022; Tan et al., 2018; Wang et al., 2018). In accord, both NRG1 and ErbB4 are risk genes of schizophrenia and major depression disorders (Howard et al., 2019; Li et al., 2025; Major Depressive Disorder Working Group of the Psychiatric Genomics Consortium, 2025; Mei and Nave, 2014). Cellular mechanisms by which the NRG1-ErbB4 signaling regulates the brain functions have been studied extensively. Manipulation of NRG1 levels or ablating or inhibiting ErbB4 disrupted the E/I balance, synaptic plasticity, LTP, sharp-wave ripples, local synchrony, and inter-regional synchrony (Agarwal et al., 2014; Chen et al., 2010b; Del Pino et al., 2013; Fazzari et al., 2010; Flames et al., 2004; Lu et al., 2014; Robinson et al., 2022; Tan et al., 2018; Wen et al., 2010; Yin et al., 2013a). However, underlying molecular mechanisms remain unclear. Increased GABA release by NRG1 and ErbB4 was shown to due to increased excitability of interneurons or local GABA release mechanism that requires nNOS activation (Li et al., 2012; Wan et al., 2024).

We speculated that in adult animals, PNN structures are not static, but rather regulated by neuronal network activity. Therefore, it would be important to investigate its roles and underlying regulatory mechanisms in the context of normal, adaptive behaviors. To this end, we subjected mice to enriched environment (EE), a physiological paradigm that is known to improve cognitive functions (Falkenberg et al., 1992; Melani et al., 2017). Remarkably, EE increased PNN levels in the prefrontal cortex in association with improved cognitive function. To investigate how EE elevates PNN levels, we performed single-nucleus RNA-seq analyses, which revealed the NRG1-ErbB4 signaling as the most significantly elevated pathway by EE from excitatory neurons to interneurons. We explored the consequences of ErbB4 mutation and chemical genetic inhibition in vivo in naïve mice and in vitro. Among the components increased by EE included Acan, a CSPG critical to PNN assembly (Giamanco et al., 2010; Rowlands et al., 2018; Suttkus et al., 2014). We therefore deleted Acan specifically in ErbB4+ interneurons and determined whether deficits caused by ErbB4 mutation could be ameliorated by ACAN expression. Results suggest that EE improves cognitive functions by enhancing PNN levels via stimulating ErbB4. This mechanism is also critical to cognitive functions in naïve mice, suggesting the dependence of PNN on ErbB4 under physiological conditions. These findings reveal a signaling pathway critical to regulate PNN levels in adult animals.

## Results

### Increased PNN levels and working memory by enriched environment

To determine whether the PNN is regulated by EE, mice were divided into two groups, one being housed in regular home cages (HC, as control) whereas the other group in enlarged cages (86×76×24 cm) that contained different sizes and shapes of tubes, a running wheel, stocks, and nest boxes (Figure 1A, designated enriched environment or EE) following a well-established protocol (Nithianantharajah and Hannan, 2006; Rampon et al., 2000; van Praag et al., 2000). Fourteen days later, mice were subjected to the Y-maze test. There was no difference in the total number of arm entries between HC and EE mice (Figure 1B-1D); however, EE mice increased spontaneous alternations in the Y-maze, compared with the HC group (Figure 1E), suggesting enhanced spatial working memory by EE. To validate EE’s effects, mice were tested for recognition memory in novel object test (Figure 1F). As illustrated in Figures 1G and 1H, both EE and HC mice exhibited preference for the novel Object B. However, EE mice showed a higher discrimination index, compared with HC mice (Figure 1I). In contrast, the total travel distance of EE and HC mice in open field arena was similar (Figure S1A and S1B), indicative of similar motor activity of the groups. Together, these results indicate that EE increased cognitive function in mice, in agreement with earlier reports (Falkenberg et al., 1992; Melani et al., 2017).

**Figure 1.**
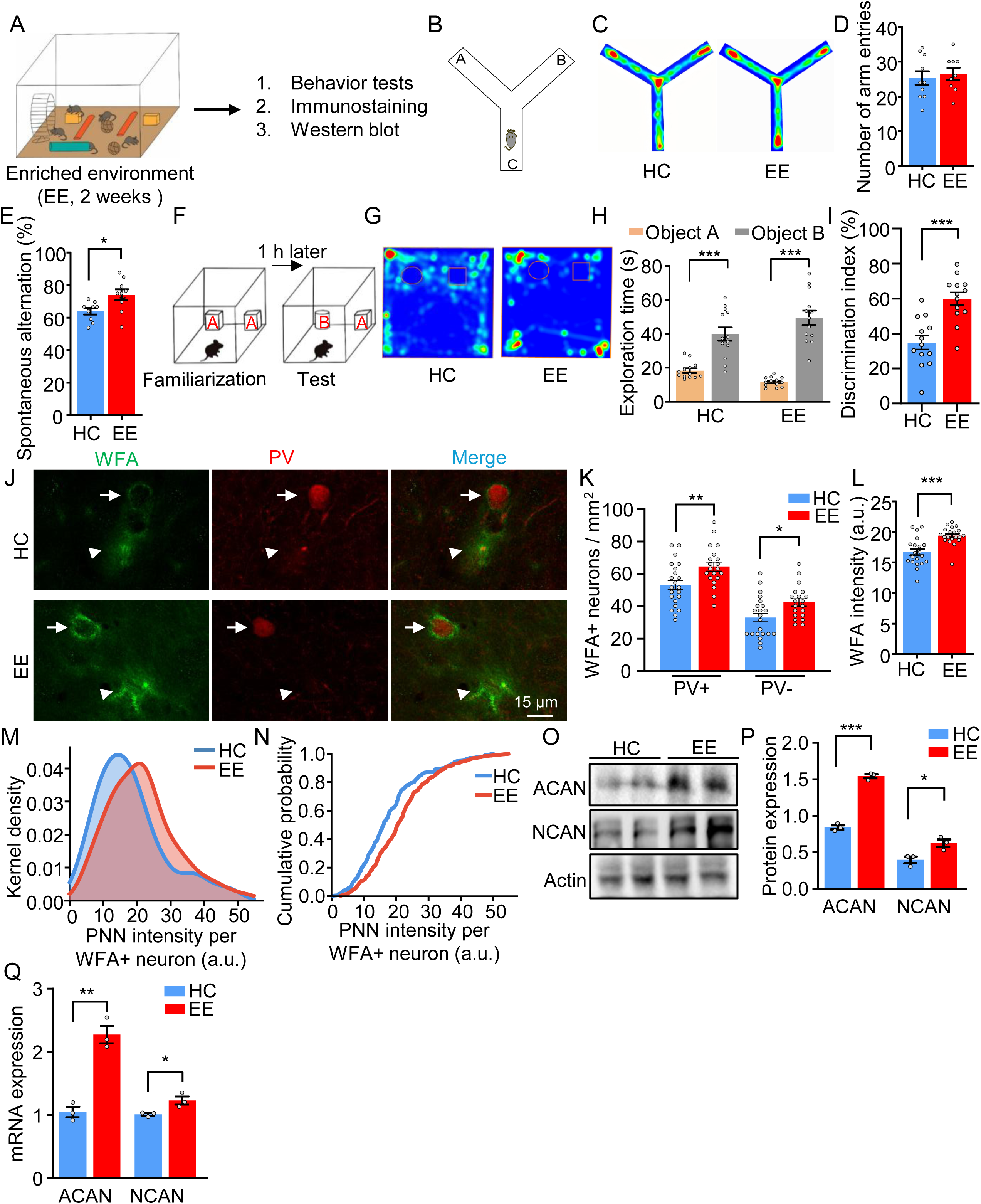
Enriched Environment Promotes PNN Expression in the PFC and Enhances Cognitive function. **(A)** Schematic of the EE (2 weeks) paradigm and experimental design. **(B-E)** Increased spontaneous alternation in the Y-maze test in EE mice. B, schematic of the Y-maze test. C, representative heatmaps of mouse activity during the Y-maze test in HC and EE groups. D, no significant difference in the number of arm entries between groups. E, spontaneous alternation in EE and HC mice. *n* = 10 mice per group. *p < 0.05; unpaired Student’s *t* test. **(F-I)** Increased discrimination index in the NOR test in EE mice. F, schematic of the NOR test. G, representative heatmaps of mouse activity during the NOR test in HC and EE groups. H, exploration time for objects A and B. I, discrimination index in HC and EE mice. *n* = 13 mice per group. \*\*\**p* < 0.001; two-way ANOVA (H) and unpaired Student’s *t* test (I). **(J-L)** Increased PNN expression in the PFC of EE mice. J, representative enlarged images of WFA labeling and PV immunostaining in the PFC of HC and EE mice; arrows indicate PV+/WFA+ neurons and arrowheads indicate PV-/WFA+ neurons. Scale bar, 15 μm. K, density of WFA+ neurons that are PV+ or PV-. L, WFA intensity in HC and EE mice. *n* = 20 images from 3 mice per group. \**p* < 0.05, \*\**p* < 0.01, \*\*\**p* < 0.001; unpaired Student’s *t* test (K and L). **(M and N)** Kernel density estimation (M) and cumulative distribution (N) plots of WFA intensity per WFA+ neuron in HC (blue) and EE (red) mice. Kolmogorov–Smirnov test, D = 0.21, *p* < 0.001. *n* = 469 neurons from 3 mice in HC; *n* = 316 neurons from 3 mice in EE. **(O and P)** Increased ACAN and NCAN protein levels in EE mice. O, representative immunoblots of ACAN and NCAN in the PFC of HC and EE mice. P, quantification of the immunoblots in (O). *n* = 3 mice per group. \**p* < 0.05, \*\*\**p* < 0.001; unpaired Student’s *t* test. **(Q)** Increased mRNA levels of *Acan* and *Ncan* in EE mice. *n* = 3 mice per group. \**p* < 0.05, \*\**p* < 0.01, unpaired Student’s *t* test. Data are presented as mean ± SEM. See also Figure S1.

To determine whether PNN levels are changed by EE, PFC sections were stained for WFA (Wisteria floribunda agglutinin) that selectively binds the glycosaminoglycan (GAG) sugar side chains of PNN glycoproteins and thus has been served as a marker of PNN (Härtig et al., 1992; Pizzorusso et al., 2002). As shown in Figures S1C and S1D, WFA+ cells were detected in layers 5/6 and L2/3 where neurons receive inputs from multiple brain regions, but rarely in layer 1 where neurons are sparse (Meyer et al., 2011). In the cortex, PNN was mostly in proximity of interneurons (Celio and Blumcke, 1994; Fawcett et al., 2019; Härtig et al., 1992; Kwok et al., 2011). In agreement, a majority (∼60%) of WFA+ cells co-stained with anti-PV antibodies (Figure 1J) although WFA was also detected in PV-interneurons. The numbers of WFA+ cells were increased in EE group (Figure 1K); in addition, the WFA intensity was higher in PFC sections of EE mice than HC mice (Figure 1L). Kernel density estimation revealed a rightward shift in the PNN intensity distribution in EE mice compared to HC controls, indicating an overall increase in PNN intensity across the population (Figure 1M). Consistently, cumulative distribution analysis showed that the EE group exhibited a significant redistribution toward higher intensity values (Figure 1N) (Kolmogorov–Smirnov test, D = 0.21, p < 0.001), suggesting a population-wide enhancement of PNN in EE mice. Note that PV intensity distributions (both Kernel and cumulative) were similar between EE mice and HC mice, indicative of the specificity of WFA intensity shift (Figure S1E and S1F, Kolmogorov–Smirnov test, D = 0.096, p = 0.341). To validate the results of morphological studies, we analyzed the expression of PNN components by RT-PCR and western blot analysis. As shown in Figure 1O-1Q, ACAN and NCAN were increased at both mRNA and protein levels in the EE group, compared with the HC group. These results indicate that EE increased PNN expression in association with improved cognitive function and suggest a potential role of PNN in cognitive function.

### Identification of NRG1-ErbB4 signaling for PNN levels

To investigate how EE increased the PNN, we took an unbiased approach to analyze the transcriptome alterations by EE. PFC tissues of HC and EE mice were subjected to single-nucleus RNA sequencing (snRNA-seq; Figure 2A). Analysis of 10,514 nuclei from the HC group and 9,525 nuclei from the EE group identified six major cell types based on well-characterized markers (Yao et al., 2023): excitatory neurons, inhibitory neurons, oligodendrocytes, oligodendrocyte precursor cells, microglia, and astrocytes (Figure 2B). The cell type classification and distributions of EE and HC mice were similar (Figure S2A and S2B), suggesting that EE had little effect on cell-type populations. We calculated PNN gene module scores of each nucleus based on the core PNN structural components including Acan, brevican, Ncan, verican, Hapln1, Hapln4, Tnr, Ptprz1. Interestingly, the cumulative fraction exhibited a right-shift in the EE group compared with HC (Figure 2C; Kolmogorov–Smirnov test, D = 0.068, p = 3.69 × 10⁻³). When analyzed by cell types, PNN module scores were increased in both excitatory and GABAergic neurons of EE mice, however with a greater increase in the latter (i.e., GABAergic) than the former group (Δ ≈ 0.031 vs 0.008) (Figure 2D), in agreement with increased WFA staining around interneurons (Figure 1L). The scores were not changed for astrocytes, microglia, oligodendrocytes or oligodendrocyte precursor cells, suggesting that EE may not have an effect on the expression of PNN components in these cells. The scores in excitatory neurons were higher in EE mice, compared with HC, suggesting increased expression of PNN genes in excitatory neurons although they could not be stained by WFA.

**Figure 2.**
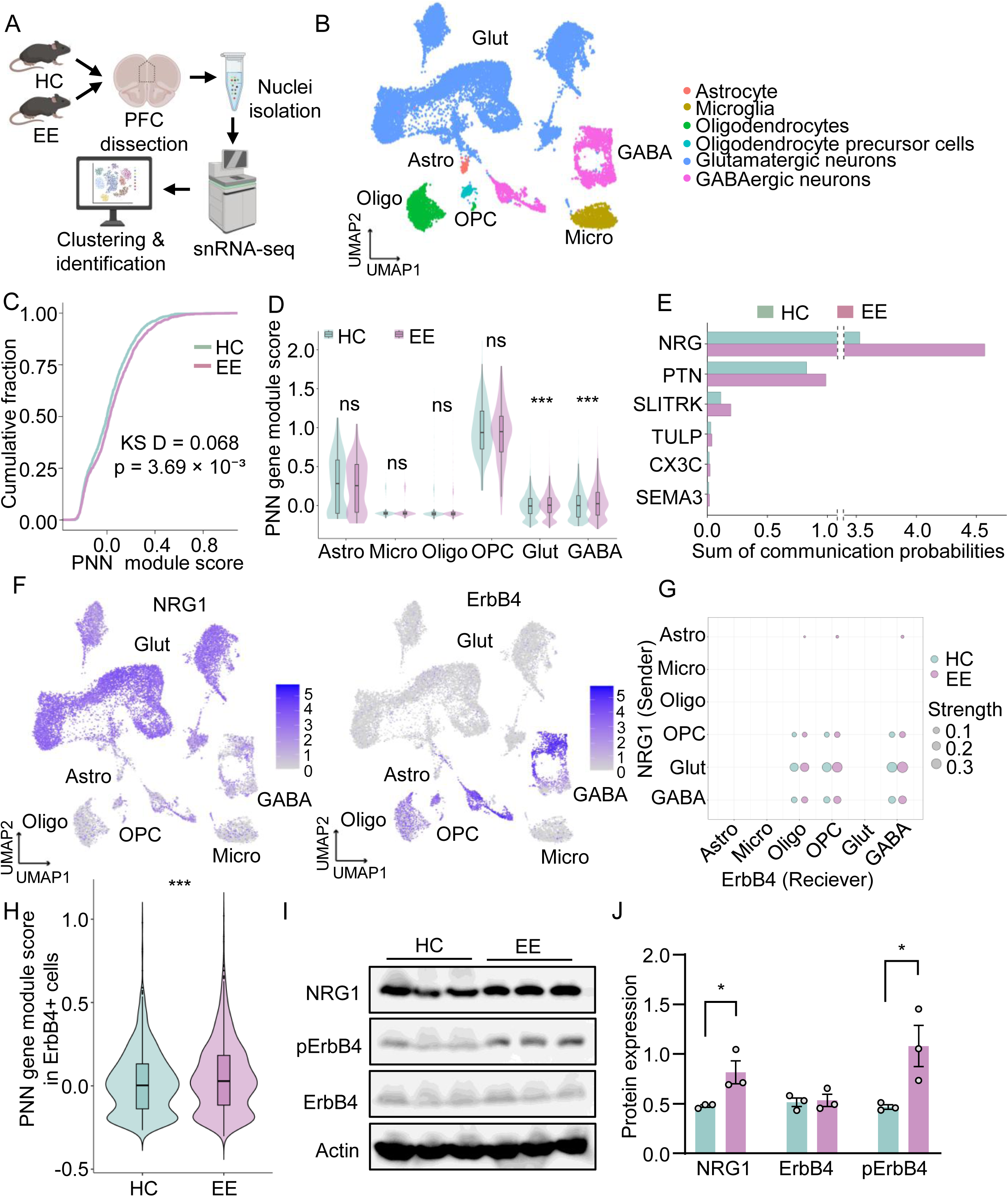
Single-Nucleus RNA Sequencing Reveals Enhanced NRG1–ErbB4 Signaling in the PFC after EE. **(A)** Schematic of the experimental workflow for snRNA-seq. **(B)** UMAP plot showing the clustering and annotation of cell populations identified by snRNA-seq. **(C and D)** Higher PNN gene module scores after EE. C, CDFs of PNN gene module scores in nuclei from HC and EE groups; Kolmogorov–Smirnov test, *D* = 0.068, *p* = 3.69 × 10^-3^. D, PNN module scores in different cell types in HC and EE groups. \*\*\**p* < 0.001, Wilcoxon rank-sum test. **(E)** Inferred CCC pathways with higher communication probabilities in the EE group. **(F)** UMAP visualization of *Nrg1* and *Erbb4* transcript expression across annotated cell types. **(G)** Comparison of inferred NRG1–ErbB4 communication strength between HC and EE groups. **(H)** Violin plots showing PNN module scores in *Erbb4*+ nuclei of HC and EE groups.\*\*\**p* < 0.001, Wilcoxon rank-sum test. **(I and J)** Increased NRG1 and pErbB4 levels in EE mice. I, representative immunoblots of NRG1, total ErbB4, and pErbB4 in the PFC of HC and EE mice. J, quantification of the immunoblots in (I). *n* = 3 mice per group. \**p* < 0.05, unpaired Student’s *t* test. Bar graphs show mean ± SEM. See also Figure S2.

PNN is often associated with interneurons and its levels are likely regulated by the activity of the cortical network (Devienne et al., 2021; McRae et al., 2007; Pizzorusso et al., 2002). To identify the intercellular communications that were altered by EE, we quantified condition-dependent changes in the number and strength of cell–cell interactions by CellChat analysis of snRNA-seq data (Jin et al., 2021). As shown in Figure S2C, the number of inferred cell-cell interactions was similar between EE and HC mice; in contrast, there was a marked increase in the strength of inferred cell-cell interactions in EE samples, compared with the HC group. This result suggests that EE may reshape the strength of existing communication channels without establishing new ones. To identify the pathways underlying this reorganization, we ranked the signaling pathways by their information flow among the six types of cells (Figure 2B). The information flow of a given pathway is defined as the sum of the communication probabilities (i.e., interaction strength) over sender cells–receiver cells pairs within its inferred network, thereby representing the overall signaling strength of a particular pathway across the population. As shown in Figure 2E, eight pathways were altered significantly (P < 0.05, paired Wilcoxon test): including six that were up-regulated in the EE group: Neuregulin (NRG), Pleiotrophin (PTN), SLIT and NTRK-like family (SLITRK), Tubby-like protein (TULP), CX3C, and Semaphorin 3 (SEMA3) (Figure 2E) and two that were down-regulated in EE: Protein Tyrosine Phosphatase Receptor Type (PTPR) and Fibroblast Growth Factor (FGF) (Figure S2D).

Considering that PNN is mostly detected around interneurons, the NRG signaling was noteworthy among the upregulated pathways. Earlier work and this study (Figure 2F) demonstrate that NRG1 is prominently expressed in excitatory neurons and ErbB4 is specifically in interneurons in the PFC in previous studies (Fazzari et al., 2010; Liu et al., 2011; Tan et al., 2018; Woo et al., 2007). These observations support a model that NRG1 from excitatory neurons activates ErbB4 to mediate PNN expression that is increased by EE. To test this hypothesis, we decomposed the NRG1–ErbB4 communication by cell types. As shown in Figure 2G and S2E, the NRG1 signaling was indeed directional, flowing from glutamatergic cells (as senders) to GABAergic cells (as receiver). The signaling flow strength was increased in EE, compared with HC. In addition, NRG2– and NRG3–ErbB4 signaling showed the same directionality (Figure S2F– S2I). These results are consistent with the observations that sensory deprivation reducing excitatory drive to PV interneurons delays or reduces PNN formation (McRae et al., 2007; Pizzorusso et al., 2002), whereas chemogenetic silencing of cortical excitatory neurons in adulthood reduced PNN expression (Devienne et al., 2021)

To further associate the NRG1–ErbB4 signaling to PNN, we examined the PNN module scores in ErbB4+ nuclei, which were increased in ErbB4+ nuclei of EE mice compared with HC (Figure 2H), although ErbB4 expression itself was comparable between the two groups (Figure S2J). Finally, we determined whether the NRG1–ErbB4 signaling is activated in the EE PFC by western blot analysis. As shown in Figure 2I and 2J, NRG1 was increased, and so was phosphorylated ErbB4 (pErbB4), an indicator of the activity of the receptor tyrosine kinase, in PFCs of EE mice compared with HC mice. In contrast, ErbB4 protein levels were similar between EE and HC groups. Taken together, these results suggest that the NRG1–ErbB4 signaling pathway may play a role in EE-enhanced cognitive function and PNN formation.

### ErbB4 kinase for PNN and cognitive function

To obtain genetic evidence for ErbB4 regulation of the PNN, we first explored the consequences of ErbB4 mutation. Although ErbB4 is specifically expressed in interneurons in the cortex in adult mice (Bean et al., 2014), its mutation during development could impair interneuron migration and differentiation to form dendrites, axons and synapses, which would prevent clear interpretation of results (Bernard et al., 2022; Lin et al., 2018; Ting et al., 2011; Yin et al., 2013b). Therefore, we crossed CAG::CreER mice with ErbB4^f/f^ mice, to generate CAG::CreER;ErbB4^f/f^ mice, which upon tamoxifen (Tam) treatment fail to express ErbB4. We referred Tam-treated CAG::CreER;ErbB4^f/f^ mice as B4iKO mice (Wang et al., 2018) (Figure 3A). ErbB4 mRNA and protein levels were reduced in the PFC of B4iKO mice, compared with control mice (i.e., Veh-treated CAG::CreER;ErbB4^f/f^ mice), indicative of successful ErbB4 ablation (Figure 3E-3G) (Wang et al., 2018). Notably, the number of WFA+ cells and the WFA intensity were reduced in the PFC in ErbB4-iKO mice, compared with control mice (Figure 3B-3D and S3A). In accord, B4iKO reduced ACAN at mRNA and protein levels but had little effect on NCAN levels (Figure 3E-3G). These results demonstrate a necessary role of ErbB4 in maintaining PNN expression. Earlier works of ours and others have demonstrated that ErbB4 mutation impairs working memory (Del Pino et al., 2013; Robinson et al., 2022; Yang et al., 2026) In agreement, B4iKO mice displayed a reduction in spontaneous alternations, compared with controls, without a change in total arm entries (Figure 3H-3I). Moreover, in the novel object recognition task, unlike control mice that spent more time exploring Object B over Object A; B4iKO mice spent similar times towards both A and B objects with a reduced discrimination index (Figures 3J-3K). These results demonstrated an association of reduced PNN and impaired working and cognition memory in B4iKO mice.

**Figure 3.**
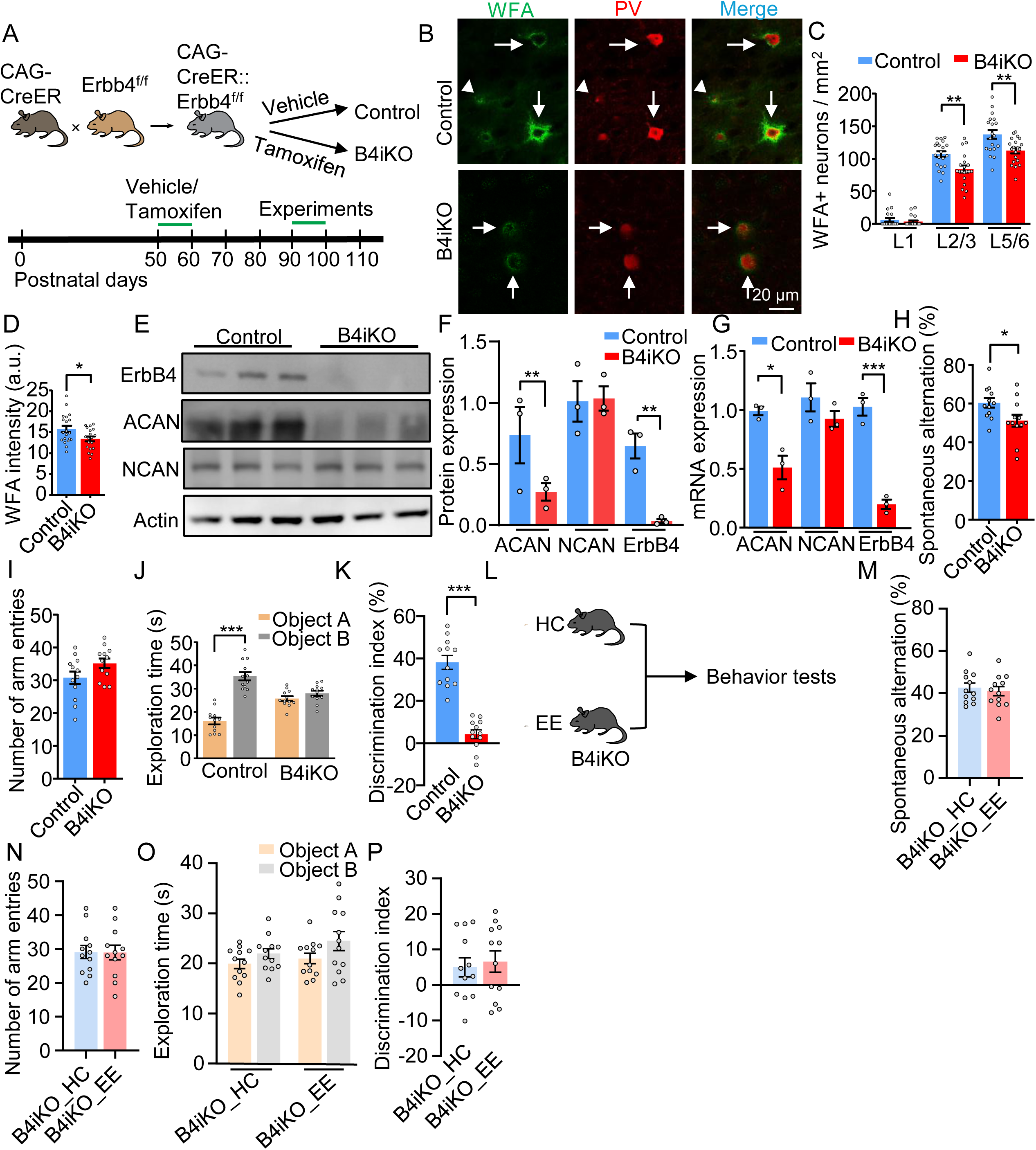
Conditional Deletion of ErbB4 Impairs PNN Expression and Abolishes EE-Induced Improvement of Cognitive function. **(A)** Schematic of tamoxifen-induced *Erbb4* deletion in CAG-CreER;Erbb4-floxed mice and the experimental timeline (B4iKO. **(B-D)** Impaired PNN expression in B4iKO mice. B, representative enlarged images of WFA labeling and PV immunostaining in the PFC of control and B4iKO mice. Scale bar, 20 μm. C, density of WFA+ neurons across PFC layers in control and B4iKO mice. D, WFA intensity in control and B4iKO mice. *n* = 20 images from 3 mice per group. \**p* < 0.05, \*\**p* < 0.01; one-way ANOVA (C) and unpaired Student’s *t* test (D). **(E and F)** Decreased ACAN and ErbB4 protein levels in B4iKO mice. E, representative immunoblots of ACAN, NCAN, and ErbB4 in control and B4iKO mice. F, quantification of the immunoblots in (E). *n* = 3 mice per group. \*\**p* < 0.01, unpaired Student’s *t* test. **(G)** *Acan*, *Ncan*, and *Erbb4* mRNA levels in control and B4iKO mice. *n* = 3 mice per group. \**p* < 0.05, \*\*\**p* < 0.001, unpaired Student’s *t* test. **(H and I)** Decreased spontaneous alternation of Y-maze test in B4iKO mice. H, spontaneous alternation in control and B4iKO mice. I, no significant difference in the number of arm entries. *n* = 12 mice per group. \**p* < 0.05; unpaired Student’s *t* test. **(J and K)** Decreased discrimination index in the NOR test in B4iKO mice. J, exploration time for objects A and B. K, discrimination index in control and B4iKO mice. *n* = 12 mice per group. \*\*\**p* < 0.001; two-way ANOVA (J) and unpaired Student’s *t* test (K). **(L)** Schematic of the experimental design for the EE intervention in B4iKO mice. **(M and N)** No significant difference in spontaneous alternation in the Y-maze test between B4iKO_HC and B4iKO_EE groups. M, spontaneous alternation. N, number of arm entries. *n* = 12 mice per group. Not significant; unpaired Student’s *t* test. **(O and P)** No significant difference in discrimination index in the NOR test between B4iKO_HC and B4iKO_EE groups. O, exploration time for objects A and B. P, discrimination index. *n* = 12 mice per group. Not significant; two-way ANOVA (O) and unpaired Student’s *t* test (P). Data are presented as mean ± SEM. See also Figure S3.

Next, we determined whether ErbB4 ablation prevents EE from enhancing cognitive function. B4iKO mice were randomly assigned to HC (B4iKO_HC) or EE (B4iKO_EE) housing for 14 days and subjected to behavioral assessments (Figure 3L). Remarkably, in the Y-maze test, spontaneous alternations and total numbers of arm entries of B4iKO_EE mice were similar to those of B4iKO_HC mice, indicating that EE was no longer able to enhance working memory (Figure 3M and 3N). Consistently, in the novel object recognition task, B4iKO_EE mice showed no preference for the novel object (Object B) over the familiar one (Object A), and their discrimination index was comparable to that of B4iKO_HC mice (Figure 3O–3P). These results demonstrate that EE was unable to improve cognitive functions Tam-treated B4iKO mice and suggest a necessary role of ErbB4 in EE’s effects.

ErbB4 is a transmembrane receptor tyrosine kinase and could interact in trans with NRG3 to promote excitatory synapses and with Slitrk3 to increase inhibitory synapses onto interneurons (Luo et al., 2021; Müller et al., 2018). These effects appeared to be independent of its kinase activity. To test whether PNN regulation by ErbB4 requires its kinase activity in vivo, we utilized ErbB4-T796G (T796G) mice in which the ATP-binding pocket of the kinase was enlarged by the T796G mutation, rendering ErbB4 selectively sensitive to the bulky inhibitor 1NM-PP1 (Tan et al., 2018) (Figure 4A). As shown in Figure 4E and 4F, phosphorylated ErbB4 in the PFC was reduced when T796G mice were treated with 1NM-PP1 (i.p. for 2 days), compared with mice treated with saline, indicating reduced ErbB4 kinase activity. However, ErbB4 protein and mRNA levels were similar between 1NM-PP1 and saline-treated mice (Figure 4E-4G). These results suggest that 1NM-PP1 inhibited ErbB4 kinase without altering ErbB4 expression. Compared with the saline control group, 1NM-PP1 treatment reduced WFA staining intensity, WFA+ neurons (Figure 4B-4D), and ACAN protein and mRNA in T796G mice, compared with mice treated with saline (Figure 4E-4G). These results suggest that ErbB4 kinase activity is required for PNN and ACAN expression. In addition, T796G mice treated with 1NM-PP1 reduced spontaneous alternation and were impaired in novel object recognition, compared with saline-treated group (Figure 4H-4K). Moreover, EE enhancement of working memory and novel object recognition was also diminished in T796G mice after 1NM-PP1 treatment (Figure 4L-4P). These results suggest that ErbB4 kinase activity is necessary for PNN expression, cognitive functions, as well as the beneficial effects of EE.

**Figure 4.**
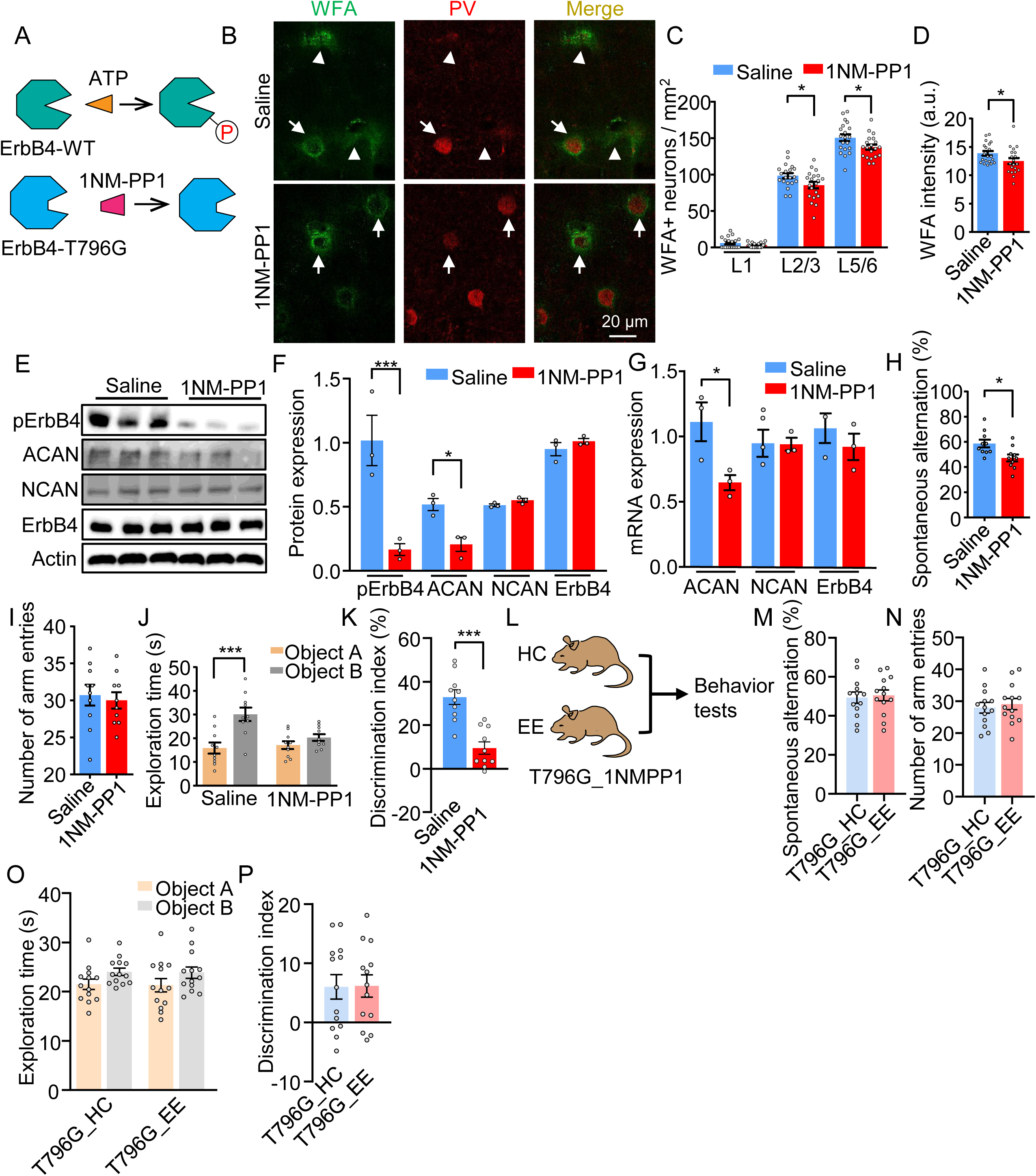
Loss of ErbB4 Kinase Activity Impairs PNN Expression and Abolishes EE-Induced Improvement of Cognitive function. **(A)** Schematic of the chemical-genetic strategy for inhibiting ErbB4-T796G kinase activity using 1NM-PP1. **(B-D)** Impaired PNN expression in 1NM-PP1-treated mice. B, representative enlarged images of WFA labeling and PV immunostaining in the PFC of saline-and 1NM-PP1-treated ErbB4-T796G mice. Scale bar, 20 μm. C, density of WFA+ neurons across PFC layers in saline- and 1NM-PP1-treated ErbB4-T796G mice. D, WFA intensity in saline- and 1NM-PP1-treated ErbB4-T796G mice. *n* = 20 images from 3 mice per group. \**p* < 0.05; one-way ANOVA (C) and unpaired Student’s *t* test (D). **(E and F)** Decreased ACAN and pErbB4 levels in 1NM-PP1-treated mice. E, representative immunoblots of ACAN, NCAN, total ErbB4, and pErbB4 in saline- and 1NM-PP1-treated mice. F, quantification of the immunoblots in (E). *n* = 3 mice per group. \**p* < 0.05, \*\*\**p* < 0.001; unpaired Student’s *t* test. **(G)** *Acan*, *Ncan*, and *Erbb4* mRNA levels in saline- and 1NM-PP1-treated mice. *n* = 3 mice per group. \**p* < 0.05, unpaired Student’s *t* test. **(H and I)** Decreased spontaneous alternation in the Y-maze test in 1NM-PP1-treated mice. H, spontaneous alternation in saline- and 1NM-PP1-treated ErbB4-T796G mice. I, no significant difference in the number of arm entries. *n* = 10 mice per group. \**p* < 0.05; unpaired Student’s *t* test. **(J and K)** Decreased discrimination index in the NOR test in 1NM-PP1-treated mice. J, exploration time for objects A and B. K, discrimination index in saline- and 1NM-PP1-treated mice. *n* = 10 mice per group. \*\*\**p* < 0.001; two-way ANOVA (J) and unpaired Student’s *t* test (K). **(L)** Schematic of the EE intervention in 1NM-PP1-treated ErbB4-T796G mice. **(M and N)** No significant difference in spontaneous alternation in the Y-maze test between HC- and EE-housed ErbB4-T796G mice treated with 1NM-PP1. M, spontaneous alternation. N, number of arm entries. *n* = 13 mice per group. Not significant; unpaired Student’s *t* test. **(O and P)** No significant difference in discrimination index in the NOR test between the groups described in (M). O, exploration time for objects A and B. P, discrimination index. *n* = 13 mice per group. Not significant; two-way ANOVA (O) and unpaired Student’s *t* test (P). Data are presented as mean ± SEM. See also Figure S4.

We reasoned that if the NRG1-ErbB4 signaling regulates PNN formation, PNN should be present in ErbB4+ interneurons. Because of the lack of reliable anti-ErbB4 antibodies, we turned to ErbB4::CreER;Ai9 (ErbB4 reporter) mice (Figure 5A). In ErbB4::CreER mice, CreER expression is under the control of ErbB4 promoter (Bean et al., 2014). Our earlier works indicate that tdTomato was specifically expressed in interneurons in the PFC, hippocampus and amygdala (Bean et al., 2014). PFC slices were prepared from Tam-treated ErbB4 reporter mice and co-stained with WFA with or without anti-PV antibodies. As observed previously, ErbB4+ interneurons were present in different layers of the cortex (Figure 5B) (Bean et al., 2014). Interestingly, 82% of WFA+ cells were positive for tdTomato, whereas 55% of PV+ cells were WFA+ (Figure 5C). Most PV+WFA+ cells expressed tdTomato. These results indicate that ErbB4 is expressed in most of WFA+ cells.

**Figure 5.**
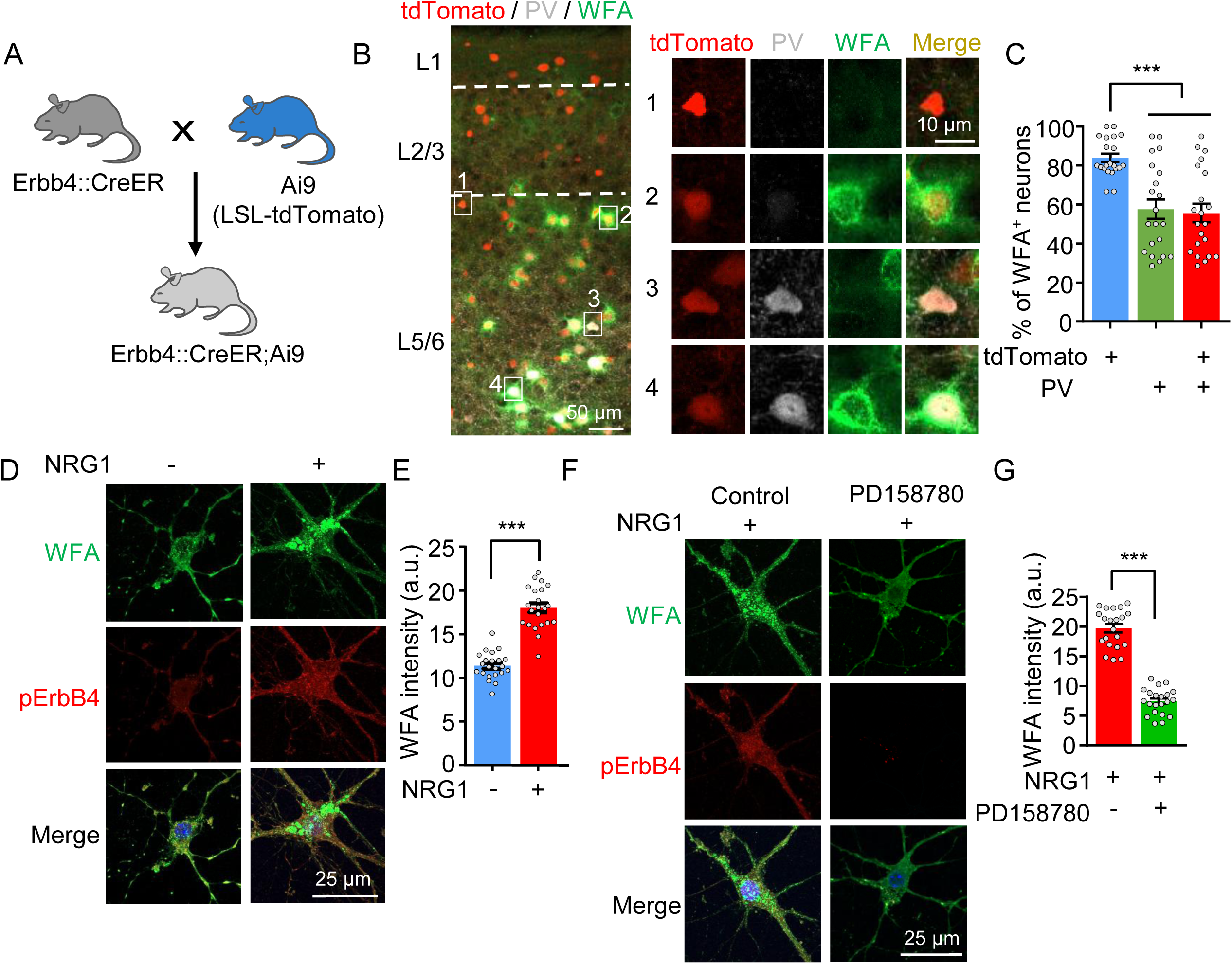
NRG1–ErbB4 Signaling Promotes PNN Expression *In Vitro* **(A-C)** A higher fraction of WFA+ neurons is tdTomato+ than PV+ in the PFC. A, schematic of the genetic strategy used to generate ErbB4-CreER;Ai9 reporter mice. B, representative images of tdTomato fluorescence, PV immunostaining, and WFA labeling in the PFC of ErbB4-CreER;Ai9 mice. Scale bars, 50 μm (overview) and 10 μm (enlarged images). C, fractions of tdTomato+, PV+, and tdTomato+/PV+ neurons among WFA+ neurons. *n* = 21 images from 3 mice. \*\*\**p* < 0.001, one-way ANOVA. **(D and E)** NRG1 increases PNN expression in cultured neurons. D, representative images of WFA labeling (green), pErbB4 immunostaining (red), and DAPI counterstaining (blue) in primary cultured neurons with or without NRG1 treatment. Scale bar, 25 μm. E, WFA intensity following NRG1 treatment. *n* = 22 neurons per group. \*\*\**p* < 0.001, unpaired Student’s *t* test. **(F and G)** PD158780 blocks the NRG1-induced increase in PNN expression. F, representative images of WFA labeling (green), pErbB4 immunostaining (red), and DAPI counterstaining (blue) in NRG1-treated primary neurons in the presence or absence of PD158780. Scale bar, 25 μm. G, WFA intensity in the NRG1 and NRG1 + PD158780 groups. *n* = 21 neurons per group. \*\*\**p* < 0.001, unpaired Student’s *t* test. Data are presented as mean ± SEM.

The above observations from studies of genetically manipulated mice suggest that PNN levels and cognitive functions require ErbB4 and its kinase activity. To demonstrate that NRG1 acts directly on interneurons to stimulate PNN expression, cortical neurons were isolated from mouse embryos, cultured for 11 days in vitro (DIV11), treated with NRG1 for 24 h, and co-stained with WFA and antibody against pErbB4. As expected, NRG1 treatment increased pErbB4 level. Importantly, NRG1 also increased the WFA staining intensity of cultured neurons (Figure 5D and 5E). Notably, the effect of NRG1 was reduced by PD158780, an inhibitor of ErbB4, compared with the control (Figure 5F and 5G). Taken together, these in vitro results suggest that NRG1 may be able to increase PNN formation by activating ErbB4 in interneurons.

Taken together, both in vivo and in vitro results provide evidence that the NRG1-ErbB4 signaling may play a role in maintaining PNN and cognitive functions and in EE enhancement of cognitive functions likely by upregulating the PNN.

### ACAN in ErbB4+ interneurons for PNN levels and glutamatergic transmission

ACAN is a core component of PNN structure whose distribution pattern closely overlaps with WFA-labeled PNN at both the cellular and tissue levels (Matthews et al., 2002; McRae et al., 2010). Its loss-of-function leads to pronounced reduction of PNN in the cortex and hippocampus (Grødem et al., 2025; Rowlands et al., 2018), in contrast to rather mild PNN deficits by other CSPG knockdowns or knockouts (Favuzzi et al., 2017; Schmidt et al., 2020). We showed that EE-induced as well as basal PNN were reduced by genetic ablation or inhibition of ErbB4, which was associated with reduced ACAN, but not NCAN (Figure 3E-3G and 4E-4G). These observations suggest ErbB4-dependent regulation of PNN levels and cognitive functions by controlling ACAN expression. In support of this hypothesis is that 95% of ACAN+ cells in the PFC were positive for ErbB4 (Figure 6A-6C and S5A). Similarly, RNA fluorescence in situ hybridization indicated ACAN was expressed in ErbB4+ neurons, regardless whether they were positive or not for PV (Figure 6C).

**Figure 6.**
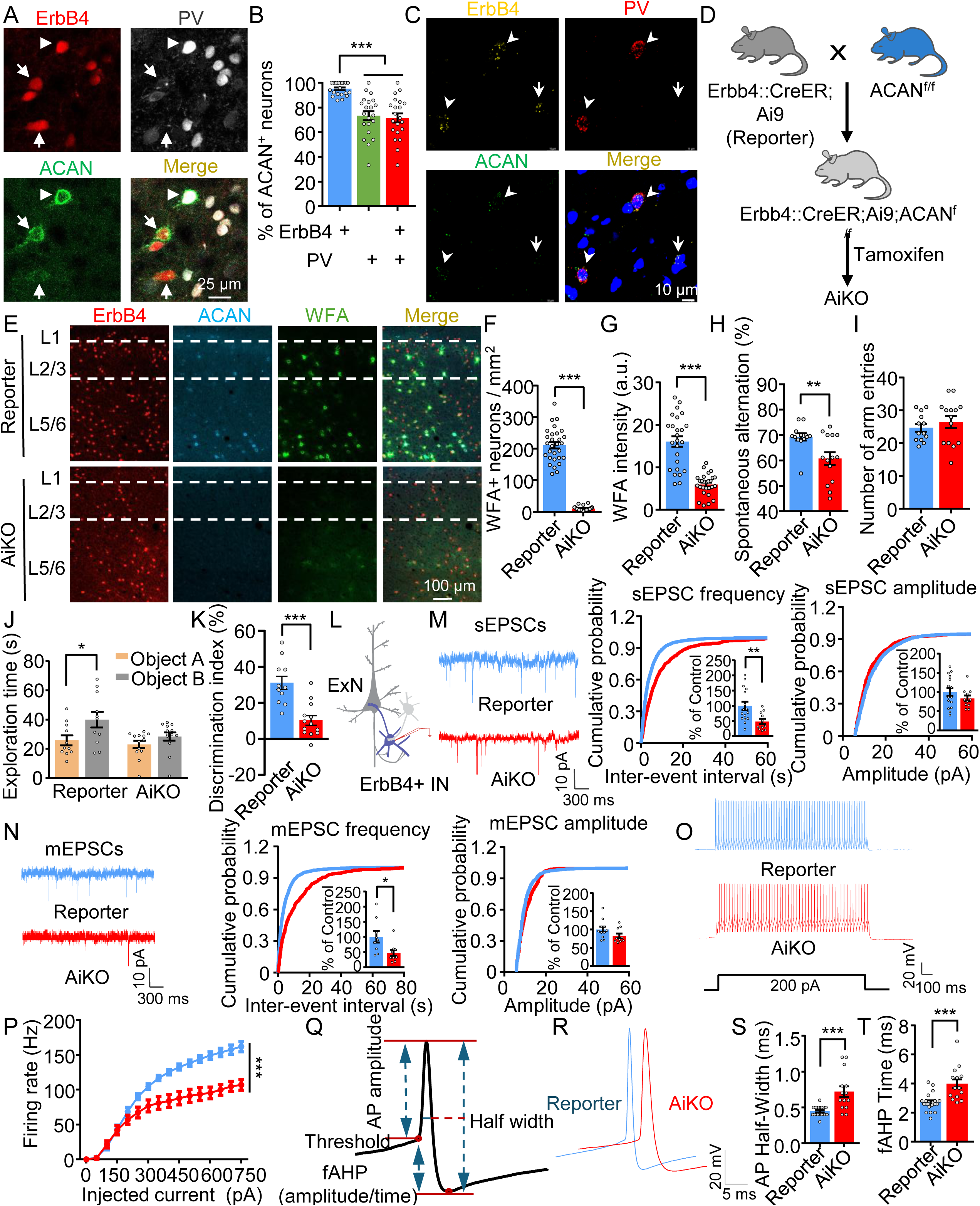
Conditional Deletion of ACAN in ErbB4+ Interneurons Impairs PNN Expression, Excitatory Synaptic Transmission, Intrinsic Excitability, and Cognitive function. **(A and B)** A higher fraction of ACAN+ neurons is ErbB4-reporter+ than PV+ in the PFC. A, representative images of PV and ACAN immunostaining together with ErbB4 reporter fluorescence in the PFC of reporter mice; arrows indicate ErbB4-reporter+/ACAN+/PV-neurons and arrowheads indicate ErbB4-reporter+/ACAN+/PV+ neurons. Scale bar, 25 μm. B, fractions of ErbB4-reporter+, PV+, and ErbB4-reporter+/PV+ neurons among ACAN+ neurons. *n* = 21 images from 3 mice. \*\*\**p* < 0.001, one-way ANOVA. **(C)** Representative RNA amplification-based single-molecule in situ hybridization images of *Erbb4*, *Pvalb*, and *Acan* transcripts in the PFC. Arrows indicate *Erbb4*+/*Acan*+/*Pvalb*-neurons; arrowheads indicate *Erbb4*+/*Acan*+/*Pvalb*+ neurons. Scale bar, 10 μm. **(D)** Schematic of the genetic strategy used to generate ErbB4-CreER;Ai9;*Acan*^f/f^ (AiKO) mice. **(E-G)** Impaired PNN expression in AiKO mice. E, representative images of WFA labeling, ACAN immunostaining, and ErbB4 reporter fluorescence in the PFC of reporter and AiKO mice. Scale bar, 100 μm. F, density of WFA+ neurons; *n* = 29 images from 4 reporter mice and 26 images from 4 AiKO mice. G, WFA intensity; *n* = 25 images from 4 reporter mice and 27 images from 4 AiKO mice. \*\*\**p* < 0.001, unpaired Student’s *t* test. **(H and I)** Decreased spontaneous alternation in the Y-maze test in AiKO mice. H, spontaneous alternation in reporter and AiKO mice. I, no significant difference in the number of arm entries. *n* = 13 reporter mice; *n* = 14 AiKO mice. \*\**p* < 0.01; unpaired Student’s *t* test. **(J and K)** Decreased discrimination index in the NOR test in AiKO mice. J, exploration time for objects A and B. K, discrimination index in reporter and AiKO mice. *n* = 11 reporter mice; *n* = 13 AiKO mice. \**p* < 0.05, \*\*\**p* < 0.001; two-way ANOVA (J) and unpaired Student’s *t* test (K). **(L)** Schematic of whole-cell patch-clamp recording from ErbB4-targeted interneurons. **(M and N)** Decreased excitatory synaptic input onto ErbB4+ interneurons in AiKO mice. M, sEPSCs; *n* = 15 neurons from 3 reporter mice and 11 neurons from 3 AiKO mice. N, mEPSCs; *n* = 9 neurons from 3 mice per group. For each, representative traces (left), CDFs of inter-event intervals and amplitudes (right), and normalized frequency and amplitude measurements (insets) in reporter and AiKO mice. Scale bars, 10 pA, 300 ms. \*\**p* < 0.01 (M) and \**p* < 0.05 (N) for frequency, unpaired Student’s *t* test. **(O and P)** Decreased firing rate of ErbB4+ interneurons in AiKO mice. O, representative traces of APs in reporter and AiKO mice. Scale bars, 20 mV, 100 ms. P, current injection–induced APs. *n* = 16 neurons from 3 reporter mice; *n* = 17 neurons from 3 AiKO mice. \*\*\**p* < 0.001, one-way ANOVA. **(Q-T)** Broadened AP waveform and prolonged fAHP in AiKO interneurons. Q, schematic defining AP threshold, amplitude, half-width, and fAHP measurements. R, representative single APs from reporter and AiKO interneurons. Scale bars, 20 mV, 5 ms. S, AP half-width. T, fAHP duration. *n* = 14 neurons from 3 reporter mice; *n* = 20 neurons from 3 AiKO mice. \*\*\**p* < 0.001, unpaired Student’s *t* test. Data are presented as mean ± SEM. See also Figures S5–S7.

To test this hypothesis, we ablated ACAN specifically in ErbB4+ neurons by crossing floxed ACAN mice with ErbB4-reporter mice (Figure 5A) to generate ErbB4::CreERT2; Ai9; ACAN^f/f^ mice (Figure 6D and S5B). Resulting mice were treated with Tam, which enabled not only ACAN ablation (Figure S5C-S5E), but also labeling ErbB4+ interneurons (Figure 6E). Tam-treated ErbB4::CreERT2; Ai9; ACAN^f/f^ mice were thus referred to as AiKO mice. Nissl staining revealed no gross anatomical abnormalities in the brains of AiKO mice (Figure S5F). However, WFA staining intensity and the number of WFA+ cells were nearly abolished in the PFC of AiKO mice (Figure 6E–6G). These findings suggest that PNN in the PFC require ACAN expression in ErbB4+ interneurons. In accord with the notion that PNN regulates cognitive functions, AiKO mice exhibited a significant reduction in spontaneous alternation behavior in the Y-maze, without changing arm entries, compared with control mice (Figure 6H-6I). Their novel object recognition was also compromised, with reduced discrimination index (Figure 6J-6K). These results suggest that ACAN from ErbB4+ interneurons is critically involved in cognitive function.

To explore the mechanisms by which PNN regulates cognitive function, we assessed synaptic transmission onto ErbB4+ interneurons by recording excitatory and inhibitory postsynaptic currents (EPSCs and IPSCs) (Figure 6L). Notably, the frequency of spontaneous EPSC (sEPSCs) was reduced in AiKO mice although sEPSC amplitudes remained unchanged (Figure 6M). In contrast, neither the frequency nor the amplitude of sIPSCs differed between AiKO and control mice (Figure S6A). Spontaneous synaptic events comprise both action potential (AP)–dependent and -independent components. We therefore examined miniature EPSCs (mEPSCs) which are generated by spontaneous vesicle release. As shown in Figure 6N, mEPSC frequency, but not amplitude, was reduced in AiKO mice, compared with control mice. By contrast, miniature IPSCs (mIPSCs) showed no differences in either frequency or amplitude between AiKO and control mice (Figure S6B). In addition to reduced synaptic inputs, AiKO also reduced the firing rate of ErbB4+ interneurons (Figure 6O and 6P). Both action potential half-width and fast afterhyperpolarization (fAHP) time were increased in AiKO mice, compared with control mice (Figure 6Q-6T). No significant differences were observed in input resistance (Figure S7A), resting membrane potential (Figure S7B), fAHP amplitude (Figure S7C), action potential threshold (Figure S7D), or action potential amplitude (Figure S7E). These results suggest that ACAN in ErbB4+ interneurons is necessary for maintaining excitatory synaptic inputs and AP firing and reveal novel functions of ACAN.

Given the widespread expression of ErbB4 throughout the brain (Bean et al., 2014), the cognitive deficits in AiKO mice may arise from ACAN ablation in various brain regions (Figure 6H-6K). Therefore, we next sought to selectively suppress ACAN expression in ErbB4+ interneurons in the PFC. To this end, three short-hairpin RNAs (shAcan) targeting mouse Acan mRNA were generated (Figure S8A) (Favuzzi et al., 2017). As shown in Figure S8B, shAcan2 was most efficient to suppress ACAN expression in cells co-transfected with HA-tagged ACAN and CAG-Cre plasmids. Resulting shAcan AAV was injected into the PFC of ErbB4-reporter mice (Figure S8C–S8D), which were treated with Tam and characterized one month after. The expression of the EGFP indicated Cre-dependent expression of AAV. As shown in Figure S8E, ACAN expression and WFA staining were largely reduced in EGFP-positive regions, confirming the effective knockdown of ACAN and associated PNN structures. Notably, ACAN knockdown in the PFC phenocopied AiKO mice in behavioral deficits including diminished working memory and novel object recognition (Figure S8F-S8I) and in neurotransmission impairment including reduced mEPSC frequency and AP firing (Figure S9 and S10). These results demonstrate that ACAN produced by ErbB4+ interneurons in the PFC is required for normal neurotransmission and working memory and novel object recognition.

### ACAN co-condensates with BCAN and NCAN

Despite the important roles of the PNN in neural development and synaptic plasticity (McRae et al., 2007; Pizzorusso et al., 2002), little is known of molecular mechanisms of its assembly. Our data that the PNN was almost abolished in AiKO mice, suggesting that ACAN may play a critical role in PNN formation. ACAN consists of a long, anionic chondroitin sulfate attachment region that is predicted to be intrinsically disordered, a feature shared by many proteins capable of liquid-liquid phase separation (LLPS) (Banani et al., 2017; Molliex et al., 2015). Condensates have been implicated in the assembly of pre- and post-synaptic apparatus (Milovanovic et al., 2018; Wu et al., 2019; Zeng et al., 2016). Therefore, we determined whether purified ACAN was able to undergo LLPS to form condensates. As shown in Figure 7A-7C and S11A, ACAN readily formed micron-scale condensates were readily detectable within 5 min of incubation. The condensates formed by BCAN were fewer albeit being similar in size with ACAN condensates. However, condensates formed by NCAN were sparce and irregular in shape, despite after extended incubation (Figure 7B and 7C). FRAP analysis indicated that the condensates were able to recover after bleaching (Figure 7D– 7E), suggesting dynamic exchange of ACAN and BCAN in and out of the condensates. FRAP analysis of NCAN condensates was difficult due to its small size. These results demonstrate that ACAN is most capable of spontaneously forming condensates, followed by BCAN and NCAN.

**Figure 7.**
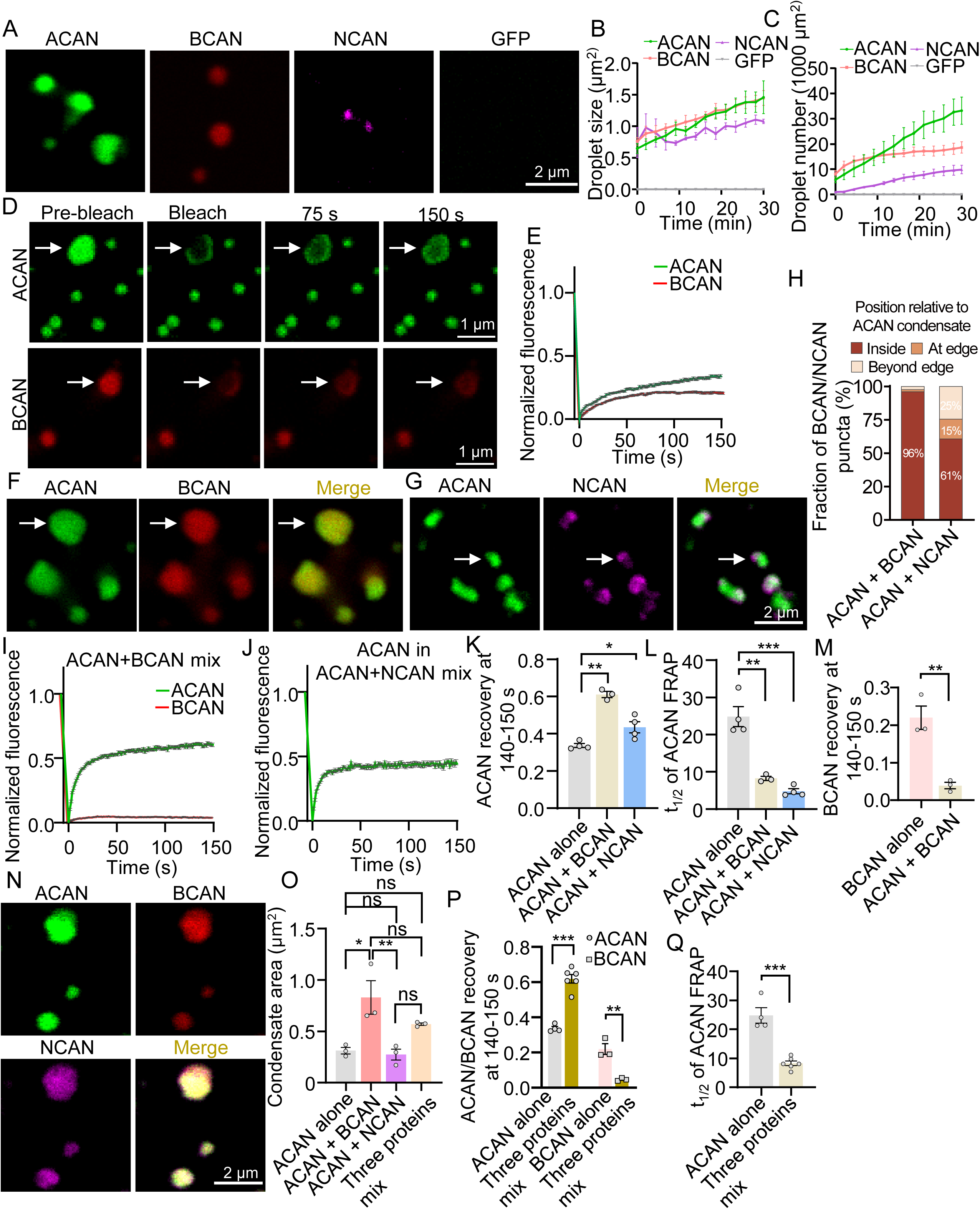
Condensate Formation and Spatial Organization of ACAN, BCAN, and NCAN. **(A-C)** ACAN, BCAN, and NCAN form condensates. A, representative fluorescence images of ACAN (green), BCAN (red), and NCAN (magenta) condensates, and the GFP control. Scale bar, 2 μm. B, droplet area over time. C, droplet number per 1,000 μm^2^ over time. *n* = 3 experimental repeats per condition and time point. **(D and E)** ACAN and BCAN condensates show partial fluorescence recovery after photobleaching. D, representative FRAP images of ACAN and BCAN condensates before bleaching, immediately after bleaching, and at 75 and 150 s after bleaching; arrows indicate the condensates subjected to photobleaching. Scale bars, 1 μm. E, normalized FRAP curves for ACAN (green) and BCAN (red) over 150 s. *n* = 45 ROIs from 4 experimental repeats in ACAN; *n* = 29 ROIs from 3 experimental repeats in BCAN. **(F-H)** BCAN and NCAN are differentially recruited into ACAN condensates. F, representative fluorescence images of ACAN + BCAN condensates. G, representative fluorescence images of ACAN + NCAN condensates; ACAN, green; BCAN, red; NCAN, magenta. Arrows indicate representative ACAN condensates containing BCAN or NCAN signals. Scale bar, 2 μm. H, distribution of BCAN and NCAN puncta inside, at the edge of, or beyond the edge of ACAN condensates; bars show the mean percentage of the same puncta as in Figure S11E. **(I-M)** Co-condensation alters the FRAP dynamics of ACAN and BCAN. I, normalized FRAP curves of ACAN and BCAN in ACAN + BCAN condensates. J, normalized FRAP curves of ACAN in ACAN + NCAN condensates. K, ACAN recovery, measured as the mean normalized fluorescence at 140–150 s after bleaching. L, half-time of ACAN fluorescence recovery (*t*_1/2_), obtained by fitting a single exponential to the mean FRAP curve of each experimental repeat. M, BCAN recovery at 140–150 s after bleaching. *n* = 45 ROIs from 4 experimental repeats in ACAN alone; *n* = 27 (ACAN) and 36 (BCAN) ROIs from 3 experimental repeats in ACAN + BCAN; *n* = 36 ROIs from 4 experimental repeats in ACAN + NCAN; *n* = 29 ROIs from 3 experimental repeats in BCAN alone. \**p* < 0.05, \*\**p* < 0.01, \*\*\**p* < 0.001; one-way ANOVA with Dunnett’s post hoc test (K and L) and unpaired Student’s *t* test (M). **(N and O)** Condensate area in the presence of BCAN, NCAN, or both. N, representative fluorescence images of ACAN + BCAN + NCAN condensates; ACAN, green; BCAN, red; NCAN, magenta. Scale bar, 2 μm. O, condensate area in ACAN alone, ACAN + BCAN, ACAN + NCAN, and ACAN + BCAN + NCAN mixtures. Each dot represents one independent preparation (median condensate area per field, averaged over 6, 7, 11 and 6 fields, respectively; n = 3 preparations; 216, 316, 658 and 322 condensates). \**p* < 0.05, \*\**p* < 0.01; ns, not significant. Two-way ANOVA with Tukey’s post hoc test. Comparisons are indicated by brackets. **(P and Q)** FRAP dynamics of ACAN and BCAN in ACAN + BCAN + NCAN condensates. P, ACAN and BCAN recovery at 140–150 s after bleaching in the three-protein mixture, compared with ACAN alone and BCAN alone, respectively. Q, *t*_1/2_ of ACAN fluorescence recovery in ACAN alone and the three-protein mixture. *n* = 49 (ACAN) and 26 (BCAN) ROIs from 6 and 3 experimental repeats, respectively, in the three-protein mixture; ACAN alone and BCAN alone data are the same as in (K)–(M). \*\**p* < 0.01, \*\*\**p* < 0.001; unpaired Student’s *t* test. For (O), each dot represents one independent preparation (mean of its fields of view); n = 3 preparations per group. For (K)–(M), (P) and (Q), each dot represents one experimental repeat (mean of its ROIs). Data are presented as mean ± SEM. See also Figure S11.

Next, we determined whether the three glycans were able to co-condensate with one another or together. As shown in Figure 7F and S11B, the ACAN-BCAN co-condensates were readily detectable, mostly rounded, similar to those formed by individual proteins. While BCAN florescence was evenly distributed in the co-condensates, NCAN seemed to be enriched in subdomain that often localized at edges of the co-condensates (Figure 7F-7H and S11C-S11E). Interestingly, after photobleaching of ACAN-BCAN co-condensates, the fluorescence of ACAN, but not BCAN, was able to recover (Figure 7I, 7K-7M and S11F). Notably, ACAN recovery was faster and to a higher level for ACAN-BCAN co-condensates than in condensates formed by ACAN alone (Figure 7K, 7L and S11F). On the other hand, the co-condensates formed by ACAN and NCAN appeared less homogenous in fluorescence distribution, as if NCAN condensates attached to those of ACAN. Apparent subdomain of NCAN were readily detectable. Nevertheless, ACAN fluorescence also recovered faster and to a higher level than in condensates formed by ACAN alone (Figure 7J–7L and S11G). These results suggest that ACAN may act as a core scaffold protein to maintain PNN stability.

Finally, we examined whether ACAN, BCAN and NCAN were able to form co-condensates when mixed together. As shown in Figure 7N and S11H, co-condensates formed by the three proteoglycans were well-rounded, similar to those formed by ACAN and BCAN (Figure 7F and S11B). This finding suggests that in the presence of both ACAN and BCAN, the ability of NCAN to form irregularly-shaped condensates was suppressed. While ACAN and BCAN appeared to be homogeneously distributed in the co-condensates, NCAN’s distribution was uneven in some co-condensates (Figure 7N and S11I). Furthermore, the size of the co-condensates was similar to those formed by ACAN and BCAN (Figure 7O). Interestingly, after photobleaching, ACAN recovered faster and to a higher level in the co-condensates than in ACAN condensates, but BCAN recovery remained low (Figure 7P–7Q and S11J). These results suggest that the proteoglycans have unique properties in liquid-liquid phase separation that could be altered by other proteoglycans in the co-condensates. Being more dynamic, ACAN condensates may serve as an initial scaffold for PNN formation.

### Necessity of ACAN in ErbB4+ interneurons for PNN expression and cognitive function

The LLPS results reminded us that experience should be unable to build new PNN without ACAN, even when the upstream NRG1–ErbB4 signal remains intact. Two observations made this prediction testable: EE increased both PNN levels and cognitive performance in control mice (Figure 1), and deleting Acan from ErbB4+ interneurons lowered PNN levels and impaired cognition at baseline (Figure 6). We therefore asked whether EE could still enhance PNN expression and cognitive function when ACAN was absent from these neurons. AiKO mice were randomly assigned to EE or HC cages (referred to as AiKO_EE and AiKO_HC, respectively) and examined 14 days later (Figure 8A). WFA staining in AiKO_HC mice was low (Figure 8B), in agreement with Figure 6E; notably, there was no difference in WFA staining between AiKO_HC and AiKO_EE, suggesting that EE was unable to increase PNN levels (Figure 8B). Unlike naïve mice whose cognitive function could be increased by EE (Figure 1B-1I), in AiKO mice, such effects were diminished (Figure 8C–8F). The firing rate, AP half-wave and fAHP time were similar between AiKO_HC and AiKO-EE mice, suggesting they remained impaired in AiKO mice (Figure 8G-8N). These results support the notion that EE enhancement of cognitive function may require ACAN expression in ErbB4+ interneurons.

**Figure 8.**
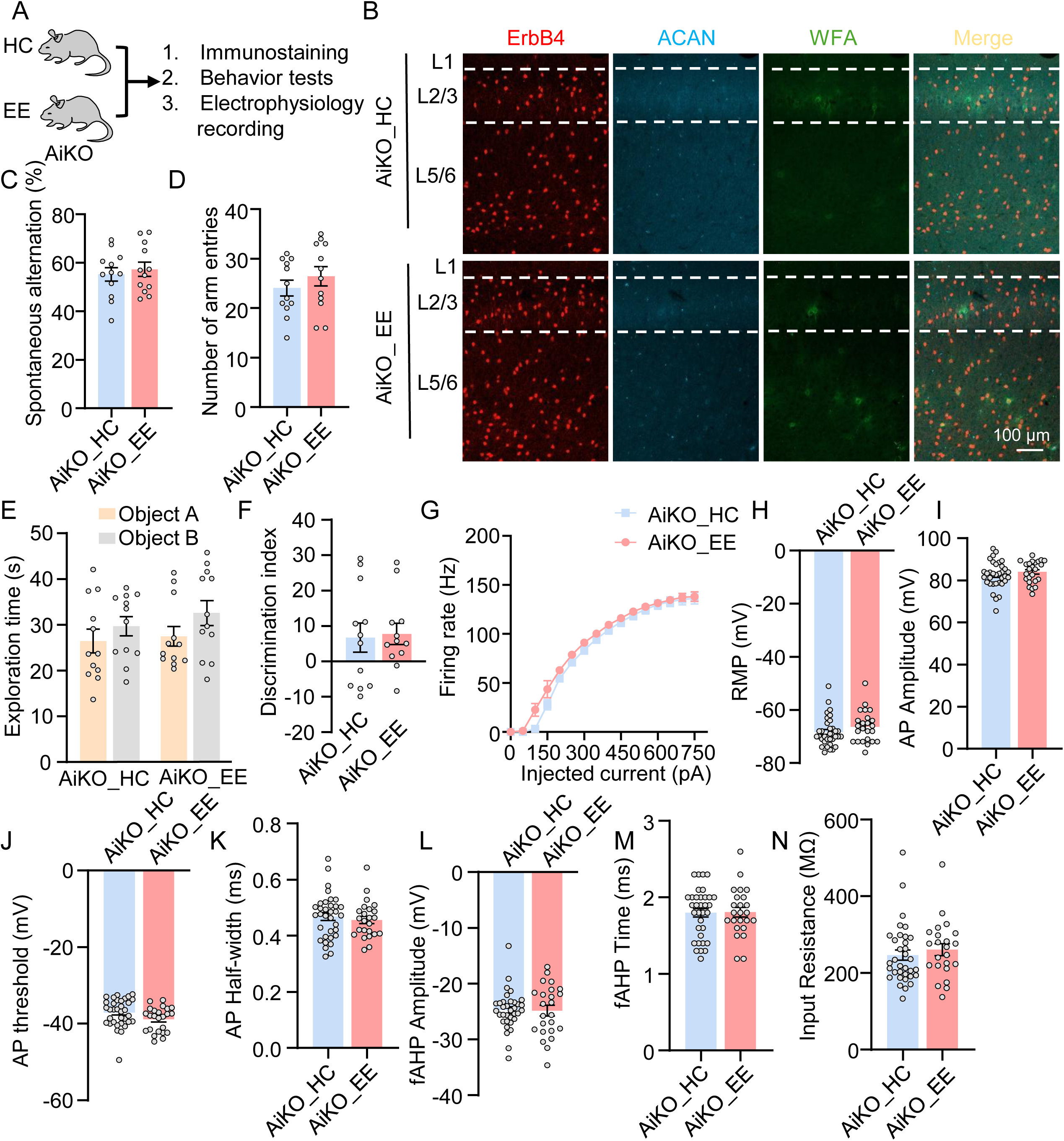
EE Fails to Rescue PNN Deficits, Cognitive Impairment, and Intrinsic Property Changes in ACAN Conditional Knockout Mice. **(A)** Schematic of the experimental design. **(B)** Representative images of WFA labeling, ACAN immunostaining, and ErbB4 reporter fluorescence in the PFC of AiKO_HC and AiKO_EE mice. Scale bar, 100 μm. **(C and D)** No significant difference in spontaneous alternation between AiKO_HC and AiKO_EE groups. C, spontaneous alternation. D, number of arm entries. *n* = 12 mice per group. **(E and F)** No significant difference in discrimination index in the NOR test between AiKO_HC and AiKO_EE groups. E, exploration time for objects A and B. F, discrimination index. *n* = 12 mice per group. **(G)** No significant difference in firing rates between AiKO_HC and AiKO_EE groups. *n* = 17 neurons from 3 AiKO_HC mice; *n* = 13 neurons from 3 AiKO_EE mice. **(H-N)** No significant difference in intrinsic membrane properties between AiKO_HC and AiKO_EE interneurons. H, RMP. I, AP amplitude. J, AP threshold. K, AP half-width. L, fAHP amplitude. M, fAHP duration. N, input resistance. *n* = 34 neurons from 3 AiKO_HC mice; *n* = 24 neurons from 3 AiKO_EE mice. Data are presented as mean ± SEM. No significant differences were detected. Statistical tests: two-way ANOVA for (E) and unpaired Student’s *t* tests for the other quantified panels.

We reasoned if ACAN acts as an effect of ErbB4, ACAN overexpression should be able to rescue the deficits caused by ErbB4 deficiency. To test this, we generated a Cre-dependent lentiviral vector for ACAN overexpression (Figure 9A). ACAN could be detected in the condition medium and lysates of CAG-Cre co-transfected HEK293T cells (Figure S12A). Resulting lentivirus were stereotaxically injected into the PFC of B4iKO mice that did not express ErbB4 in the brain (Figure 9B-9C). One month after injection, the PFC was isolated and analyzed for ACAN expression. As shown in Figure 9D and S12B, ACAN-3xHA was detected in PV+ interneurons of injected mice. Notably, WFA+ neurons were increased in L2/3 and L5/6, but not L1, of ACAN-OE mice, compared with mice injected with 3xHA virus (Figures 9E), with increased PNN intensity (Figure 9F). ACAN-3xHA was also detected by western blot in the PFC of ACAN-OE mice, which remained negative for ErbB4 (Figure 9G and 9H). Remarkably, spontaneous alternation was increased in ACAN-OE mice, compared with mice injected with 3xHA virus (Figures 9I and 9J). In addition, ACAN-OE mice showed increased preference for the novel object with increased discrimination index, compared with the control mice (Figures 9K-9L). These results suggest that ACAN overexpression in the PFC was able to rescue the PNN expression and behavioral deficits due to ErbB4 mutation, suggesting ACAN as a critical mediator of the NRG1-ErbB4 signaling for PNN expression and cognitive function.

**Figure 9.**
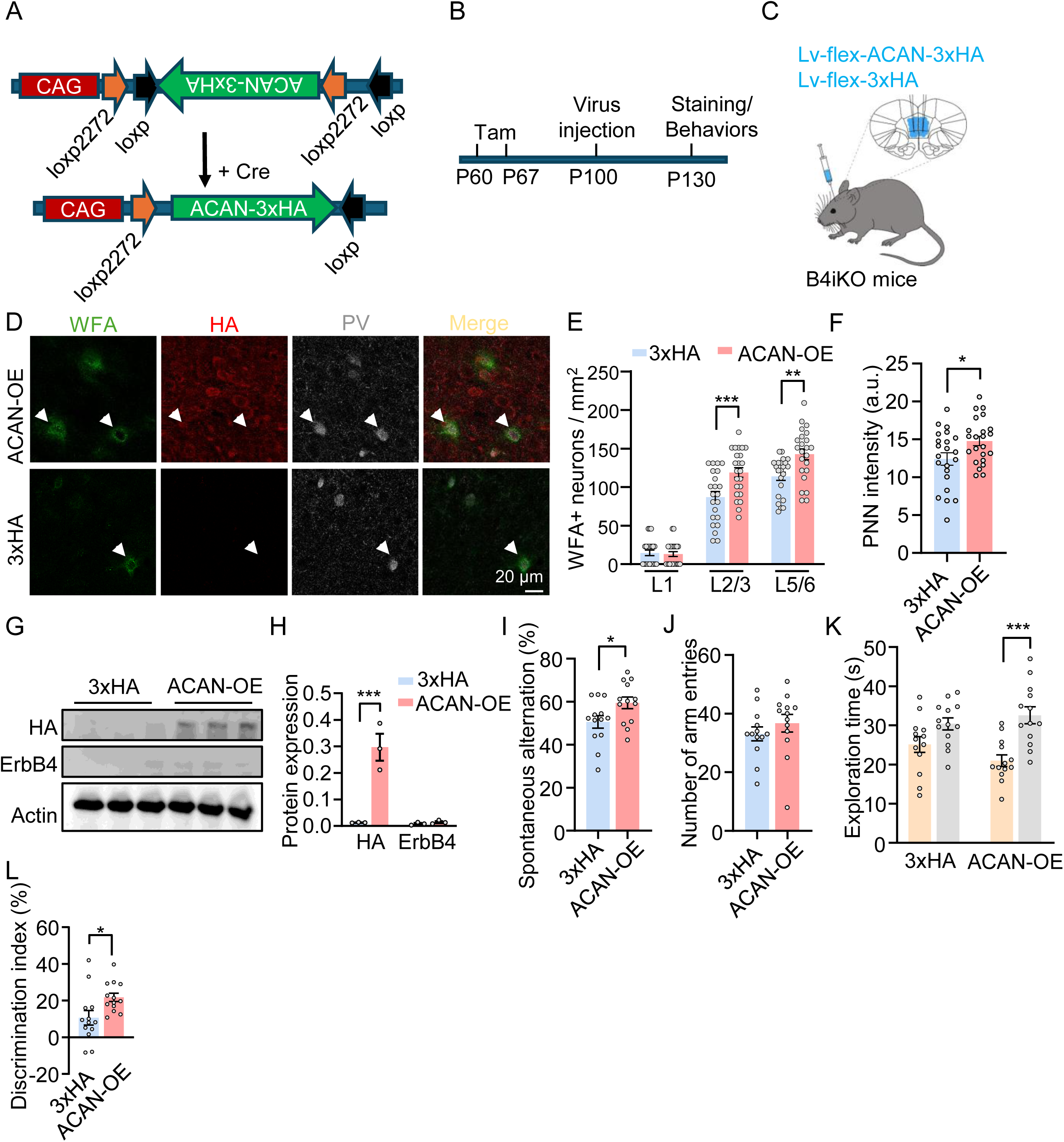
ACAN Overexpression in B4iKO Mice Rescues PNN Expression and Cognitive function. **(A-C)** Strategy for Cre-dependent ACAN overexpression in the PFC of B4iKO mice. A, schematic of the FLEX-ACAN-3×HA construct and the Cre-dependent strategy used to express ACAN-3×HA (ACAN-OE). B, experimental timeline. C, lentivirus injection and the targeted PFC injection sites. **(D-F)** ACAN overexpression increases PNN expression in B4iKO mice. D, representative enlarged images of WFA labeling and PV and HA immunostaining in the PFC of B4iKO mice following injection of the control or ACAN-OE lentivirus; arrowheads indicate PV+/WFA+ neurons. Scale bar, 20 μm. E, density of WFA+ neurons across PFC layers. F, WFA intensity. *n* = 22 images from 3 mice in the 3xHA group; *n* = 23 images from 3 mice in the ACAN-OE group. \**p* < 0.05, \*\**p* < 0.01, \*\*\**p* < 0.001; two-way ANOVA (E) and unpaired Student’s *t* test (F). **(G and H)** Lentiviral ACAN-3×HA expression in the PFC of B4iKO mice. G, representative immunoblots of HA-tagged ACAN and ErbB4 in the PFC of B4iKO mice. H, quantification of the immunoblots in (G). *n* = 3 mice per group. \*\*\**p* < 0.001, unpaired Student’s *t* test. **(I and J)** Increased spontaneous alternation in the Y-maze test in the ACAN-OE group. I, Y-maze spontaneous alternation in the 3xHA and ACAN-OE groups. J, no significant difference in the number of arm entries. *n* = 13 mice per group. \**p* < 0.05; unpaired Student’s *t* test. **(K and L)** Increased discrimination index in the NOR test in the ACAN-OE group. K, exploration time for objects A and B. L, discrimination index in the 3xHA and ACAN-OE groups. *n* = 13 mice per group. \**p* < 0.05, \*\*\**p* < 0.001; two-way ANOVA (K) and unpaired Student’s *t* test (L). Data are presented as mean ± SEM. See also Figure S12.

## Discussion

Our study reveals a signaling pathway that promotes PNN expression and its critical role in experience-enhanced cognitive function. We showed that PFC WFA staining and PNN components like ACAN were increased in EE mice, in association with improved cognitive function, compared with HC mice (Figure 1). Moreover, unbiased CellChat analysis of DEGs between EE and HC PFCs identified the NRG1-ErbB4 signaling as a potential pathway to promote PNN expression by EE. This hypothesis is supported by the following pieces of evidence. First, in the PFC, NRG1 is mostly expressed in excitatory neurons whereas ErbB4 is present exclusively in interneurons (Mei and Xiong, 2008) (Figure 2F). Second, both NRG1 levels and pErbB4, an indicator of kinase activity, were increased in EE mice, compared with HC mice (Figure 2I and 2J). Third, B4iKO and chemical genetic inhibition of ErbB4 kinase prevented EE from enhancing cognitive function and PNN levels (Figure 3L-3P and 4L-4P). Fourth, NRG1 stimulation increased PNN expression in cultured interneurons in a manner that requires ErbB4 kinase activity (Figure 5). In accord, these phenotypes were reproduced by specifical ablation of ACAN in ErbB4+ interneurons (Figure 6). Finally, we showed that re-expression of ACAN in the PFC was able to rescue the deficits of PNN and cognitive functions of ErbB4 mutation. These results provide compelling evidence that the NRG1-ErbB4 signaling is required for PNN expression under physiological conditions and is utilized by EE to promote cognitive functions via increasing PNN.

EE has been shown to optimize the E/I balance and shape theta and gamma oscillations in the PFC (Alemany-González et al., 2022; Huang et al., 2021; Sale et al., 2007), which are essential for coordinating information transfer between distant brain regions during memory encoding, consolidation, and retrieval (Jones and Wilson, 2005; Maingret et al., 2016; Place et al., 2016; Siapas et al., 2005). EE could increase the expression of BDNF, which via MAPKs increases LTP and synaptic proteins such as synaptophysin and PSD-95 in the PFC (Falkenberg et al., 1992; Nithianantharajah et al., 2004; Novkovic et al., 2015). The beneficial effects of EE also require the activation of metabotropic glutamate receptor mGluR5, which initiates mTOR signaling cascades (Hullinger et al., 2015). In addition, EE could induce epigenetic changes and activate transcription factors such as AP-1 and NPAS4 to produce long-lasting effects (Alaiz-Noya et al., 2025; Bloodgood et al., 2013). Our study demonstrates that the NRG1-ErbB4 signaling could serve as a mediator of EE-induced PNN expression in the PFC, highlighting a previously unappreciated pathway to regulate the extracellular matrix by experience. Intriguingly, inducible ablation of ErbB4 or chemical genetic inhibition of ErbB4 kinase by 1NM-PP1 reduced basal PNN levels and cognitive functions (Figure 3B-3K and 4B-4K), suggesting that brain functions require basal activity of the NRG1-ErbB4 pathway. This notion agrees with earlier findings that this pathway is required for maintaining the excitability and GABAergic transmission of interneurons (Fazzari et al., 2010; Li et al., 2012; Lu et al., 2014; Tan et al., 2012; Woo et al., 2007).

PNN has been viewed traditionally as a “brake” on plasticity due to its role in closing critical periods (Pizzorusso et al., 2002). Our findings suggest that the PNN or its components may be not merely structural stabilizers of the extracellular matrix but rather act as dynamic regulatory elements that gate experience-regulated cognitive function. The correlation between increased WFA staining and ACAN expression in EE mice and their improved cognitive function establishes a direct link between PNN integrity and higher-order brain function (Figure 1). Conversely, this relationship was reinforced by the cognitive deficits observed in mice when ErbB4 or ACAN are reduced in an inducible manner (Figure 3 and 6), demonstrating that PNN impairment is sufficient to hamper cognitive performance. Notably, ErbB4 and ACAN deficiency in adulthood prevents EE from enhancing cognitive functions (Figure 3 and 6). These results suggest that first, the increase in PNN in EE mice is not an epiphenomenon, but a necessary step for EE to exert its effects on adult cognition. Second, PNN remain dynamic in adulthood and are actively involved in the mechanisms of synaptic plasticity underlying cognitive functions regulated by EE. Whether PNN can be remodeled by other brain functions is worthy testing.

By forming a specialized lattice around interneurons, PNN have been shown to stabilize their firing patterns (Balmer, 2016; Favuzzi et al., 2017; Lensjo et al., 2017), which are essential for the temporal coordination of neuronal ensembles (Cobb et al., 1995; Pouille and Scanziani, 2001). When ErbB4+ interneurons were specifically depleted off ACAN, their firing rates were reduced, with increased action potential half-width and fast afterhyperpolarization (fAHP) time (Figure 6S-6T). In addition, the synaptic transmission of these neurons was impaired (Figure 6M-6N). We propose that the EE-induced increase in PNN enhances the inhibitory activity in the PFC to optimize the signal-to-noise ratio in PFC networks for the execution of cognitive tasks. Our data suggests that PNN serves as a critical interface between environmental experience and cognitive output. The rescue of both PNN and cognitive deficits by re-expression of ACAN (Figure 9), a key PNN component, provides causal evidence for this regulatory role.

The NRG1-ErbB4 pathway is known for its roles in promoting events critical to GABAergic circuit assembly including interneuron migration, differentiation, and formatting synapses (Mei and Nave, 2014; Mei and Xiong, 2008; Rico and Marín, 2011). In adult brains, ErbB4 deficiency impairs GABA release and synaptic plasticity (Tan et al., 2018; Woo et al., 2007) and various brain functions including working memory, attention, social novelty and fear conditioning (Bi et al., 2015; Del Pino et al., 2013; Lu et al., 2014; Robinson et al., 2022; Tan et al., 2018; Wang et al., 2018). Underlying cellular mechanisms including maintaining the E/I balance, synaptic plasticity, LTP, sharp-wave ripples, local synchrony, and inter-regional synchrony (Agarwal et al., 2014; Batista-Brito et al., 2023; Chen et al., 2010a; Del Pino et al., 2013; Fazzari et al., 2010; Lu et al., 2014; Robinson et al., 2022; Sun et al., 2016; Tan et al., 2018; Wen et al., 2010; Yin et al., 2013a). This study provides evidence that the NRG1-ErbB4 signaling may serve as an activity-dependent regulator of the extracellular matrix in the adult brain. First, unbiased CellChat analysis of DEGs between EE and HC PFCs identified the NRG1-ErbB4 signaling as a potential pathway to promote PNN expression by EE. Indeed, NRG1 levels were increased in the PFC of EE mice; so was pErbB4 although its protein levels remained unchanged (Figure 2I and 2J). Second, ErbB4 ablation or inhibition reduced PNN expression the PFC and prevented EE from improving cognitive functions (Figure 3L-3P and 4L-4P). Because ErbB4 ablation or inhibition could alter the neuronal activity, it would be important to demonstrate that NRG1 is sufficient to enhance PNN levels in a manner dependent on ErbB4 activity in a clear system. Indeed, when cultured interneurons were stimulated with NRG1, their WFA staining levels were increased; this effect was attenuated by chemical inhibition of ErbB4 (Figure 5). A parsimonious explanation of these results is that the NRG1-ErbB4 pathway may serve as a direct transducer of neuronal activity from excitatory neurons into a structural modification of the extracellular environment around interneurons. This is, to our knowledge, the first demonstration of a defined molecular pathway that actively promotes PNN formation in the adult brain.

Among the PNN components increased by EE was ACAN, which was considered a key organizer of PNN. This notion was supported by the finding that cell-specific ablation of Acan in ErbB4+ interneurons (AiKO) recapitulated the PNN deficits and cognitive impairments observed in ErbB4 mutant mice (Figure 6D-6K). This observation is consistent with the established role of ACAN as a critical structural hub for PNN assembly (Giamanco et al., 2010; Rowlands et al., 2018; Suttkus et al., 2014). ACAN is the predominant CSPG in cortical PNN, and its large size and multiple glycosaminoglycan side chains enable it to cross-link with other matrix components, including hyaluronan and link proteins. Intriguingly, we showed that ACAN underwent LLPS to form condensates, better than other forms of PNN components (Figure 8). Phase separation is increasingly recognized as a fundamental mechanism for organizing macromolecules in both intracellular and extracellular environments. By forming condensates, ACAN can create local high concentration zones that serve as nucleation sites for PNN assembly. Indeed, ACAN condensates were able to recruit BCAN that itself was unable to form large condensates, suggesting that ACAN condensates may serve as an initial scaffold, bringing together diverse CSPG components in a spatially confined manner. Importantly, LLPS offers a mechanism for rapid, reversible assembly and disassembly that is inherently responsive to local environmental conditions. Changes in ionic strength, pH, or post translational modifications can shift the LLPS equilibrium, allowing ACAN condensates to form or dissolve in response to neuronal activity.

NRG1 and ErbB4 are risk genes for schizophrenia and major depression (Howard et al., 2019; Li et al., 2025; Major Depressive Disorder Working Group of the Psychiatric Genomics Consortium, 2025; Mei and Nave, 2014); relatedly, PNN deficits were noted in post-mortem brains of psychiatric disorders (Berretta et al., 2015; Mauney et al., 2013). PNN have been shown to protect neurons from oxidative stress and excitotoxicity by buffering local ion concentrations (Cabungcal et al., 2013; Härtig et al., 1999; Morawski et al., 2004). We speculate that PNN disruption may exacerbate vulnerability of ErbB4+ interneurons, which may be metabolically demanding and particularly susceptible to oxidative damage, providing insight into pathological mechanisms of relevant brain disorders. Together, our data suggests that ACAN expression is the primary downstream effector through which NRG1-ErbB4 signaling exerts its effects on PNN integrity. This notion is supported by the rescue experiment that re-expression of ACAN in the prefrontal cortex of ErbB4 mutant mice restores both PNN levels and cognitive function.

## Resource availability

### Lead contact

Further information and requests for resources and reagents should be directed to and will be fulfilled by the lead contact, Lin Mei

### Materials availability

This study did not generate any new, unique reagents.

### Data and code availability

The reported BioProject accession for the snRNA-seq data is PRJNA1422424, with runs SRR37231793 and SRR37231792.

All original code used for condensate segmentation and quantification has been deposited at Zenodo at https://zenodo.org/records/23914857; All original code used for condensates mixture analysis and quantification has been deposited at Zenodo at https://zenodo.org/records/23915728.

Any additional information required to reanalyze the data reported in this paper is available from the lead contact upon request.

## Acknowledgements

We thank Heath Larsson Robinson (University of Montreal); Zhaoqi Dong (Capital Medical University); Xiao Yan (Chinese institute for Medical Research, Beijing); Marianne Fyhn, Gunnar Dick, Sverre Grødem, and Torkel Hafting (University of Oslo) for helpful discussions. Sequencing and bioinformatics analyses were provided by Annoroad Gene Technology (Beijing) Co., Ltd. This work was supported by the National Natural Science Foundation of China (32500839 to B.L.), the Beijing Natural Science Foundation (5254027 to B.L., QY25422 to Z.Y.M.), the China Postdoctoral Science Foundation (GZC20241105 to B.L., 2024M762175 to X.R.F.).

## Author contributions

L. M. and B. L. designed the experiments; B.L., H.L.Z., Y.M.Z and L.L.H. performed behaviors, histology and biological chemistry experiments; B.L., H.J.W. and X.F. performed the single-nucleus sequencing analysis; B.L., Z.Y.S. and W.J.Z. performed the electrophysiological recording experiments and analysis; B.L. and L.M. wrote the manuscript; All authors contributed to the data discussions.

## Materials and methods

### Animals

ErbB4::CreERT2 mice (The Jackson Laboratory, stock no. 012360) (Bean et al., 2014), inducible *Erbb4* knockout mice (B4iKO) (Wang et al., 2018), ErbB4-T796G mice (Tan et al., 2018), and Rosa26-LSL-tdTomato reporter mice (Ai9; The Jackson Laboratory, stock no. 007905), *Acan*-floxed mice (GemPharmatech, strain no. T008346) were used. ErbB4-CreER;Ai9 reporter mice were generated by crossing ErbB4::CreERT2 mice with Ai9 mice. For conditional *Acan* deletion, reporter mice were crossed with *Acan*-floxed mice to generate ErbB4-CreER;Ai9;*Acan*-flox/flox mice (AiKO). B4iKO mice carried CAG-CreER and *Erbb4*-floxed alleles.

3-4 months adult male mice were intraperitoneally administered tamoxifen at 100 mg/kg every other day over a 7-day period to induce Cre-dependent recombination; the corresponding vehicle was corn oil. Mice were housed at 22 °C under a 12-h light/12-h dark cycle with food and water available ad libitum. All animal procedures were approved by the Institutional Animal Care and Use Committee of the Chinese Institute of Medical Research, Beijing.

### Reagents and antibodies

Chemicals were purchased from Sigma-Aldrich unless otherwise specified. 1NM-PP1 was obtained from EMD Millipore (529581), and PD158780 from Calbiochem (513035). Recombinant NRG1 was prepared as described previously (Woo et al., 2007).

WFA labeling used biotin-conjugated Wisteria floribunda agglutinin (Sigma-Aldrich, L1516; 1:1,000). Primary antibodies were rabbit anti-ACAN (Sigma-Aldrich, AB1031; 1:500), sheep anti-NCAN (Invitrogen, PA5-47779; 1:1,000), rabbit anti-ErbB4 (Cell Signaling Technology, 4795; 1:1,000), rabbit anti-phospho-ErbB4 (Tyr1284; Cell Signaling Technology, 4757; 1:500), mouse anti-NRG1 (Invitrogen, MA5-12896; 1:500), rabbit anti-actin (Santa Cruz Biotechnology, sc-1616-R; 1:1,000), rabbit anti-PV (Swant, PV25; 1:1,000), mouse anti-PV (Swant, 235; 1:1,000), and rabbit anti-HA (Abcam, ab9110; 1:500) DAPI Fluoromount-G (SouthernBiotech, 0100-20, 1:5000). Fluorescent secondary antibodies were from thermos fisher, including donkey anti-rabbit Alexa Fluor 488 (A-21206), donkey anti-rabbit Alexa Fluor 568 (A10042), donkey anti-rabbit Alexa Fluor 647 (A-31573), donkey anti-mouse Alexa Fluor 647 (A-31571), donkey anti-mouse Alexa Fluor 568 (A10037), Alexa Fluor 488 conjugated streptavidin (S32354). HRP conjugated secondary antibodies were from thermos fisher, including goat anti-rabbit (31460), goat anti-mouse (31430), donkey anti-sheep (A16047).

### Environmental enrichment

Adult mice were randomly assigned to enriched environment (EE) or home cage (HC) at 12 weeks of age, balanced for litter and body weight. EE mice were housed in groups of same-sex animals in a large cage (86×76×24 cm) containing a running wheel (Ø [12] cm), plastic tunnels, a shelter, nesting material, and objects differing in shape and size. To maintain novelty, object positions were rearranged and half of the objects were replaced with new ones per week. HC mice were housed in standard cages (29 × 18 × 13 cm) with bedding and nesting material only. Both groups received the same bedding, chow and water ad libitum, and were kept on the same rack under a 12 h:12 h light–dark cycle, 22 ± 1 °C. Cages were changed weekly for both groups on the same day. Animals remained in their housing condition for 2 weeks before experiments and stayed in it throughout testing.

### Chemical-genetic inhibition of ErbB4 kinase activity

1NM-PP1 was dissolved in DMSO and diluted in sterile saline; vehicle was the identical saline–DMSO solution without drug. Adult ErbB4-T796G mice were i.p. injected with 0.1ug/g 1NM-PP1 every two hours in two days before behaviors and immunostaining. Experiments combined with EE used 2-week pumps (RWD, 1002W) with surgery performed on day 0 of EE. For surgery, mice under isoflurane anesthesia, a brain infusion cannula (RWD) was implanted into the right lateral ventricle (AP −0.4 mm, ML +1.0 mm from bregma, DV-2 mm) and connected to an osmotic minipump implanted subcutaneously between the scapulae. Mice with incompletely emptied pumps or cannula placement outside the lateral ventricle were excluded, and experimenters were blind to genotype and treatment during testing and quantification.

### Western blotting and Coomassie staining

Tissues were homogenized in PBS supplemented with protease and phosphatase inhibitors. An equal volume of 2× RIPA buffer (0.2% SDS, 1% sodium deoxycholate, and 2% Nonidet P-40 in PBS), also supplemented with inhibitors, was added, and samples were lysed for 30 min at 4 °C. Lysates were centrifuged at 12,000 × g for 15 min at 4 °C. Protein concentrations in the supernatants were determined using a BCA protein assay kit (Pierce, 23225). Samples were digested with chondroitinase ABC (chABC; 1.7 U/mg protein) for 2 h at 37 °C.

Proteins were denatured, separated by SDS-PAGE, and transferred to nitrocellulose membranes (Millipore, IPVH00010). Membranes were incubated with primary and secondary antibodies, and bands were visualized by enhanced chemiluminescence using a LI-COR imaging system (model 2800). Band intensities were quantified using ImageJ (NIH). Actin was shown as the loading control in the result figures.

### Quantitative reverse-transcription PCR

PFC tissue was dissected from individual mice, and total RNA was extracted using TRIzol (Invitrogen, 15596-026). Up to 5 μg of RNA was reverse-transcribed using a cDNA synthesis kit (Promega, A5000). Quantitative PCR was performed using the QuantiFast SYBR Green PCR Kit (Qiagen, 204054) on a StepOnePlus Real-Time PCR System (Applied Biosystems). Primer sequences (5′–3′) were: *Acan*, forward CCTGCTACTTCATCGACCCC and reverse AGATGCTGTTGACTCGAACCT; *Ncan*, forward TGCAACCACGGCTAAGCTC and reverse GGGGATAAGCAGGCAATGAC; *Erbb4*, forward CCTGCGGTCTATCCGAGAAGT and reverse ATTTAGTATTTCGGTCAGGTT; and *Gapdh*, forward AGGTCGGTGTGAACGGATTTG and reverse TGTAGACCATGTAGTTGAGGTCA.

### Cell culture

Primary cortical neurons were prepared from embryos at 18.5 days as described previously (Bean et al., 2014). Cortices were dissected and digested with 0.25% trypsin and 0.1 mg/mL DNase I for 9 min at 37 °C. Digestion was stopped by adding an equal volume of plating medium consisting of Neurobasal medium (Invitrogen, 21103049) supplemented with 10% fetal bovine serum (FBS, Gibco, A5256701). Cells were collected by centrifugation at 1100 rpm for 5 mins, resuspended in plating medium, and seeded at 6 × 10⁴ cells per well in poly-D-lysine-coated 12-well plates. After 2–4 h, plating medium was completely replaced with maintenance medium: Neurobasal medium containing 2% B-27 (Gibco, 17504044), 1% GlutaMAX (Gibco, 35050061), 50 mM glucose, and 1% penicillin–streptomycin (Gibco, 10378016). Neurons were maintained at 37 °C in 5% CO2, with half of the medium replaced every 3 days.

HEK293T cells were cultured with DMEM (Bio-Channel, BC-M-005-500mL), containing 10%FBS at incubator with 37℃, 5% CO_2_. DPBS and 0.25% trypsin (Gibco, 25200-072) were used for cell digestion.

HEK293F cells were grown in 293F Hi-exp culture medium (OPM, AC601501) at incubator with 37℃, 5% CO_2_, 120 rpm shaking conditions.

### Immunohistochemistry and cultured-neuron staining

Mice were deeply anesthetized and transcardially perfused with PBS followed by ice-cold 4% paraformaldehyde (PFA). Brains were postfixed in 4% PFA overnight at 4 °C, embedded in 2% agarose, and sectioned coronally at 40 μm using a vibratome (Leica, VT1000S). Sections were rinsed for 10 min in 0.5M TBS and blocked for 1 h at room temperature in 10% donkey serum and 0.5% Triton X-100 in TBS. Sections were incubated with primary antibodies overnight at 4 °C, washed three times with TBS, incubated with secondary antibodies for 2 h at room temperature, washed three times. Then DAPI were incubated with sections for 5 mins and washed for 2 times and mounted using AQUA-MOUNT (Lerner Laboratories, 13800).

Cultured neurons were fixed with 4% PFA in −20 °C for 15 min, washed three times with PBS, and incubated with primary antibodies overnight at 4 °C. After three washes in buffer containing 20 mM phosphate and 0.5 M NaCl, neurons were incubated with secondary antibodies for 1 h at room temperature, washed three times with PBS, and mounted on Superfrost Plus slides (Thermo fisher, 1255015).

### Nissl staining

Brain sections were mounted on glass slides, air-dried, stained with 0.1% cresyl violet for 5–10 min, rinsed briefly in distilled water, dehydrated through 70%, 95%, and 100% ethanol, and cleared in xylene. Sections were coverslipped using xylene-compatible mounting medium. Images were acquired using a VS200 slide scanner.

### RNA amplification-based single-molecule in situ hybridization

SEERNA® FISH RNA Fluorescence in Situ Hybridization Kit (Dynamic Biosystems, Suzhou, China) was used in this assay, the probes of *Erbb4* conjugated with cy5, *Pvalb* conjugated with cy3, and *Acan* conjugated with Alexa fluor 488 were customized based on mouse transcripts. Tissue sections in 10-μm thickness were first fixed in 4% (w/v) PFA for 5 minutes, and washed twice with diethyl pyrocarbonate (DEPC)-treated PBS, followed by dehydration in an ethanol series. After three washes with Wash Buffer (Dynamic Biosystems, Suzhou, China), sections were permeabilized with 0.1M HCl for 5 min in an incubator at 42°C. The sections were then incubated in order with the following solutions: target probes in hybridization mix at 42°C for 4 hours, ligation mix at 42°C for 30 minutes, splint primers in the circularization mix at 42°C for 30 minutes, amplification mix for rolling circle amplification (RCA) at 42°C for 2 hours. Slides were washed with Wash Buffer (Dynamic Biosystems, Suzhou, China) three times after each step. After that, the sections were incubated in hybridization buffer containing the label probes in a 42°C incubator for 30 min. Finally, the slides were mounted with SlowFade Gold Antifade Mountant (Thermo Fisher Scientific, Shanghai, China) medium containing 0.5 μg/mL DAPI (Sigma, Shanghai, China), which were then ready for image acquisition using the Leica confocal with 60x objective lens. For multiplex detection, equimolar amounts of target probes and label probes were used.

### Behavioral assays

Behavioral tests were performed on independent cohorts of mice. Behavior was recorded using an overhead camera and analyzed using ANY-maze (Stoelting). EthoVision (Noldus) or OBS Studio was also used for tracking or video acquisition as applicable.

#### Open field

Mice explored an empty arena (50 × 50 × 10 cm) freely for 10 min, as described previously (Wang et al., 2018). Total distance traveled was quantified.

#### Y-maze

The maze consisted of three identical 42-cm arms separated by 120°. Mice were placed in the same starting arm and explored freely for 8 min. Arm entries and consecutive three-arm sequences were recorded. Spontaneous alternation was calculated as 100 × number of consecutive triplets containing all three different arms / (total arm entries −2).

#### Novel object recognition

During familiarization, two identical objects were placed in two corners of the arena, and mice explored for 10 min before returning to their home cage. After a 1-h retention interval, one familiar object was replaced by an object differing in size, shape, and color. Mice explored the arena for 5 min, and the time spent exploring each object was quantified. The discrimination index was 100 × (Time in novel – Time in familiar) / (Time in novel + Time in familiar).

### Single-nucleus suspension preparation

Nuclear isolation was performed on ice. Frozen tissue was minced and lysed for 7 min in 1 mL NST buffer containing 0.1% NP-40, 10 mM Tris-HCl (pH 7.4), 146 mM NaCl, 1 mM CaCl2, 21 mM MgCl2, and 1 U/μL RNase inhibitor (Thermo fisher, EO0382). Cell lysis was assessed by trypan blue staining and light microscopy. Next, 1 mL ST wash buffer (10 mM Tris-HCl, 146 mM NaCl, 1 mM CaCl2, 21 mM MgCl2, and 0.01% BSA; MACS, 1000076), supplemented with 40 U/mL RNase inhibitor, was added. Suspensions were passed through a 40-μm strainer (Falcon, 352340) into a 15-mL tube; the strainer was rinsed with additional wash buffer, and filtrates were combined. Nuclei were pelleted at 500 × g for 5 min at 4 °C using a swinging-bucket rotor, resuspended in 5 mL PBS containing 1% BSA, and centrifuged again under the same conditions. The final pellet was resuspended in 100 μL PBS containing 1% BSA. Nuclear concentration and integrity were assessed by trypan blue staining and manual counting.

### Single-nucleus RNA sequencing

Libraries were prepared using the Chromium GEM-X Single Cell 3′ Kit v3.1 (10x Genomics, PN-1000268). Nuclear suspensions at 700–1,200 nuclei/μL, measured using a CellDrop FL counter, were loaded to target 10,000 nuclei per sample. Nuclei were co-encapsulated with barcoded gel beads in gel beads-in-emulsion (GEMs), followed by reverse transcription and cDNA amplification according to the manufacturer’s protocol. Reverse transcription was performed using a T100 thermal cycler (Bio-Rad) at 53 °C for 45 min and 85 °C for 5 min, followed by a 4 °C hold. cDNA concentration and quality were assessed using a Qubit fluorometer and Agilent 2100 Bioanalyzer, respectively. Libraries were sequenced by Annoroad Gene Technology on an Illumina platform at approximately 40,000 reads per nucleus.

### Single-nucleus RNA-seq processing and annotation

FASTQ files from two samples were processed using the Cell Ranger count pipeline (v7.1.0; 10x Genomics) with the mouse reference genome Mus musculus GRCm39 v1 to generate gene–barcode count matrices. Putative doublets were identified and removed using both Scrublet and DoubletFinder. Filtered count matrices were subsequently imported into Seurat (v4.1.1) (Stuart et al., 2019) in R (v4.3.1) to generate Seurat objects. Genes expressed in fewer than three cells were excluded (min. cells = 3). Cells were retained for downstream analysis if they expressed between 200 and 10,000 detected genes (nFeature_RNA) and if mitochondrial transcripts accounted for less than 20% of total UMIs (percent.mt < 0.2). Cells failing to meet these quality control criteria, as well as predicted doublets, were removed. After quality control, 9,525 cells from the EE group and 10,514 cells from the HC group were retained for downstream analyses. The EE dataset exhibited a median of 3,263 UMIs and 1,844 detected genes per cell, whereas the HC dataset showed a median of 3,112 UMIs and 1,785 detected genes per cell. Raw counts were normalized using a global-scaling normalization method with a scale factor of 10,000 per cell and log-transformed [log(count + 1)]. The top 2,000 highly variable genes (HVGs) were identified and used for downstream integration and dimensionality reduction. Data was scaled and subjected to principal component analysis (PCA) using default parameters. Significant principal components were selected based on the elbow point of the PC standard deviation plot and were subsequently used for nearest-neighbor graph construction, Louvain clustering (Traag et al., 2019), and visualization by uniform manifold approximation and projection (UMAP) (McInnes et al., 2018). The same analytical workflow was applied to identify and analyze subclusters.

### Cell–cell communication analysis

The potential CCC between cell groups were analyzed using the CellChat software (Jin et al., 2021). CellChat requires the normalized expression matrix of the sc/snRNA-seq data and the annotation of nuclei types as the inputs. Based on its built-in knowledge database of ligand-receptor interactions (LRI), CellChat calculates the strength of an L-R pair from sender cell type *i* to receiver cell type *j* (denoted as *P_ij_*) using a Hill function, specified as:

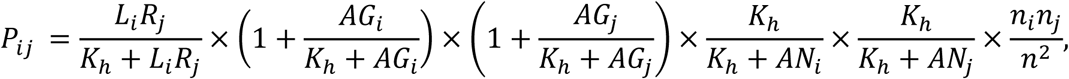

where *L_i_* and *R_j_* represents the average expression levels of the ligand and receptor in the cell group *i* and *j*, respectively. *AG* and *AN* represent the average expression of agonists and antagonists from both sender and receiver cells, whose information is also provided by the LRI database. *n_i_*, *n_j_* and *n* represent the number of sender cells, receiver cells, and all cells, respectively. *K*_ℎ_ is the parameter of the Hill function and was set to 0.5 by default.

To measure the significance level of a given LRI, CellChat performs a permutation test by randomly setting the annotation label of cells and re-calculating the LRI strength. Suppose there are *M* independent permutations in total, then the p-value can be calculated as:

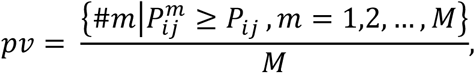

where *P^m^_ij_* denotes the calculated LRI strength of the *m*-th permutation. We applied CellChat workflow to the HC and EE samples separately and integrate the results using the mergeCellChat() function. The differentially activated LRIs were identified and visualized using the rankNet() and netVisual_diffInteraction() functions. The significance of the difference was tested by the Wilcoxon test applied on each LRI. To quantify the difference of a CCC pathway (a collection of relevant LRIs), its information flow from sender cell type *i* to receiver cell type *j* is defined as:

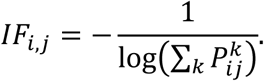

Here, *P^k^_ij_* represents the communication strength of the k-th LRI that belong to this pathway. The information of the pathways and their relevant LRIs is also included in the CellChat’s built-in knowledge base.

### Virus production

For AAV production, HEK293T cells in 15-cm dishes were co-transfected using PEI with the transfer vector (AAV-DIO-shACAN-EGFP or AAV-DIO-scramble-EGFP), pHelper, and AAV2/9 rep/cap plasmid at 10, 20, and 10 μg per dish, respectively. Cells were cultured for 5 days, and supernatants were collected. Cells were harvested and lysed in 150 mM NaCl and 50 mM Tris-HCl (pH 8.0). Lysates were treated with salt-active nuclease (SAN; ArcticZymes, 70910-202; 100 U/mL) for 30 min at 37 °C and centrifuged at 4,500 rpm for 30 min at 4 °C. Supernatants were precipitated overnight at 4 °C using PEG8000, followed by centrifugation at 4,000 × g for 30 min at 4 °C. Pellets were resuspended in SAN buffer (25 mM Tris-HCl, pH 8.5, 5 mM MgCl2, and 0.5 M NaCl) and digested with SAN for 1 h at 37 °C. Clarified material was purified by iodixanol-gradient ultracentrifugation using a Type 70 Ti rotor (Beckman Coulter) at 350,000 × g for 2 h at 18 °C. The 40% iodixanol fraction was collected and concentrated using a 100-kDa-MWCO concentrator (Thermo Fisher Scientific, 88528), aliquoted, and stored at −80 °C. Vector genome titers were measured by qPCR targeting WPRE (forward, GGCTGTTGGGCACTGACAAT; reverse, CCGAAGGGACGTAGCAGAAG). Preparations with titers ≥1 × 10¹¹ genome copies/mL were used for injection.

For lentivirus production, HEK293T cells at approximately 40%–50% confluence were transfected using PEI with FLEX-ACAN-3×HA or the FLEX-3×HA control, CMV-VSV-G, RSV-rev, and MDL packaging plasmids. Supernatant was collected 3 days after transfection, replaced with fresh medium, and collected again 3 days later. Supernatants were centrifuged at 2,000 rpm for 3 min, filtered through 0.45-μm filters, and ultracentrifuged at 20,000 rpm for 2 h at 4 °C. Pellets were resuspended in PBS, transferred to new tubes, and centrifuged with 20% sucrose at 20,000 rpm for 2 h at 4 °C. Final pellets were resuspended in 80 μL PBS, aliquoted, and stored at −80 °C.

### Stereotaxic injections

Adult mice were anesthetized with isoflurane and received bilateral PFC injections of the indicated AAV or lentivirus using a nanoliter injection pump (RWD, R-480) at 30 nL/min. The reported injection volume was 200 nL per site. Coordinates were AP - 1.90 mm, ML ± 0.30 mm from bregma, and DV + 1.70 mm target PFC. The injection needle was left in place for 5 min before slow withdrawal. Perfusion or electrophysiological recording was performed at least 1 month after injection. Injection sites were verified after experiments.

### Slice preparation and whole-cell electrophysiology

Mice were anesthetized with isoflurane and rapidly decapitated. Brains were removed into ice-cold cutting solution equilibrated with 95% O2/5% CO2 and containing (mM) 250 sucrose, 26 NaHCO3, 10 glucose, 10 MgSO4, 2 KCl, 1.3 NaH2PO4, and 0.2 CaCl2. Coronal PFC slices (300 μm) were cut using a VT1000S vibratome (Leica), incubated at 34 °C for 30 min, and allowed to recover for 1 h at 25 ± 1 °C in oxygenated artificial cerebrospinal fluid (ACSF) containing (mM) 126 NaCl, 26 NaHCO3, 10 glucose, 3 KCl, 2 CaCl2, 1.25 NaH2PO4, and 1 MgSO4. Slices were transferred to a recording chamber perfused with ACSF. Neurons were visualized with a BX51WI upright microscope (Olympus), a 40× water-immersion objective, and an infrared-sensitive CCD camera (Hamamatsu, C2400-75).

Recordings were obtained using a MultiClamp 700B amplifier, Digidata 1550B digitizer, and pCLAMP software (Molecular Devices). ErbB4-lineage neurons located at layer 5/6 were identified by tdTomato fluorescence; recorded cells were filled with biocytin and processed for post hoc WFA labeling. Signals were low-pass filtered at 2 kHz and digitized at 10 kHz. Pipette resistance was 4– 6 MΩ. The potassium-based internal solution contained (mM) 105 K-gluconate, 30 KCl, 10 HEPES, 10 phosphocreatine, 4 Mg-ATP, 0.3 Na-GTP, 0.3 EGTA, and 2.7 biocytin (pH 7.35, 285 mOsm). Series resistance was maintained below 30 MΩ.

sEPSCs were recorded at −70 mV in 20 μM bicuculline; mEPSCs were recorded with the additional presence of 1 μM tetrodotoxin (TTX). For sIPSCs, the 105 mM K-gluconate and 30 mM KCl in the internal solution were replaced with 130 mM CsCl, and pH was adjusted to 7.3 with CsOH. ACSF contained 20 μM CNQX and 50 μM DL-AP5; mIPSCs were recorded with 1 μM TTX added. Intrinsic membrane properties were measured in current-clamp mode in the presence of 20 μM CNQX, 100 μM DL-AP5, and 20 μM bicuculline. Resting membrane potential was recorded without current injection, and input resistance was calculated from voltage responses to small hyperpolarizing current steps. Depolarizing current steps were used to assess firing frequency and AP properties, including threshold, amplitude, and half-width. fAHP amplitude was measured relative to AP threshold, and fAHP latency was defined as the interval from threshold crossing to the subsequent voltage minimum. Synaptic events were detected offline and visually inspected to exclude artifacts. Event frequency and mean amplitude were calculated for each neuron, and cumulative distributions were constructed for event amplitudes and inter-event intervals.

### Recombinant protein expression and purification

Mouse ACAN, BCAN, and NCAN expression constructs were generated in a PEG2-10×His backbone with sfGFP, mKO2, or mKalama1 tags, respectively. Constructs were transfected into HEK293F cells using linear PEI (40 kDa; Polysciences, 24765-1). Sodium butyrate was added to 10 mM at 12 h after transfection. After 3 days, conditioned medium was collected, centrifuged at 12,000 rpm for 30 min at 4 °C, and filtered through a 0.45-μm filter (Millipore, HNWP04700). Filtered medium was incubated with Ni-NTA resin overnight at 4 °C. Resin was washed with five resin volumes of 500 mM NaCl, 25 mM HEPES, and 40 mM imidazole (pH 7.4). Proteins were eluted with ten resin volumes of 500 mM NaCl, 25 mM Tris, and 300 mM imidazole (pH 7.4).

### Condensate assays and fluorescence imaging

Purified proteins were diluted into imaging buffer (final concentration:10 μM) containing 150 mM NaCl, 25 mM HEPES, 6% PEG8000, and 5 mM MgCl2 (pH 7.4) at room temperature. For mixed-protein assays, components were combined at equal concentrations and diluted into the same buffer. Samples were placed in flat-bottom cavity slides (25 × 75 × 1 mm, ultra-clear optical glass), covered with coverslips, and imaged using a Leica confocal microscope with a 100× oil-immersion objective.

Time-lapse series were calibrated against a scale bar recorded in the same field of view (0.2250 µm per pixel at 512 × 512; 0.1125 µm per pixel at 1024 × 1024; field of view 115.19 × 115.19 µm = 1.327 × 10⁴ µm²) and analysed with a custom Python script (Python 3.13, scikit-image 0.26) applied identically to every series. Each frame was smoothed with a Gaussian kernel of σ = 0.225 µm for segmentation only, and a single intensity threshold, computed once from the pooled pixels of all frames of a series by Otsu’s method, was applied unchanged to every frame so that all timepoints are directly comparable. Connected components were labelled with 8-connectivity and objects below 0.20 µm² were discarded; intensities were measured on unsmoothed data after subtracting the per-frame median of sub-threshold pixels. Condensate size is reported as the median object area per frame (areas are strongly right-skewed and touching condensates segment as one object, so medians are more robust than means), and number density as the object count divided by the field-of-view area, expressed per 1,000 µm². Condensates in contact were not separated by watershed, so number density is a conservative estimate at the highest densities.

### Fluorescence recovery after photobleaching

Selected ROIs of ACAN and BCAN condensates were photobleached using 80% laser intensity. Images were acquired before bleaching, immediately afterward, and during recovery up to 150 s. Prebleach fluorescence was normalized to 100%.

### Condensate segmentation and recruitment analysis

#### Image analysis

Confocal images were exported from Leica .lif files as raw single-channel TIFFs with Bio-Formats in Fiji (Linkert et al., 2010; Schindelin et al., 2012) and analyzed with custom Python scripts based on scikit-image and SciPy (van der Walt et al., 2014; Virtanen et al., 2020). The offset between the ACAN and client channels was measured in each image by sub-pixel phase cross-correlation (Guizar-Sicairos et al., 2008) and corrected before analysis. ACAN condensates and BCAN or NCAN puncta were segmented separately by Gaussian smoothing (σ = 1 pixel) and Otsu thresholding (Otsu, 1979), and objects smaller than 9 pixels were discarded. Intensities were measured on unsmoothed images, and the dilute phase was defined as all pixels more than 3 pixels away from any segmented object. For display, brightness and contrast were adjusted linearly and identically across compared images.

#### Co-condensation and condensate size

Co-condensation and condensate size were quantified automatically in every image with custom Python. The scripts recruitment_quant.py and shape_analysis.py using the regionprops function of scikit-image to measure the area and intensity of each object. An ACAN condensate was scored BCAN- or NCAN-positive when its mean client intensity exceeded the dilute-phase mean + 3 SD. A client punctum was scored as inside an ACAN condensate when at least 50% of its area overlapped the ACAN mask, at the edge when it overlapped less but touched the mask after 1-pixel dilation, and beyond the edge otherwise. Condensate area was calculated from the ACAN mask, and the median area of each field was used.

### Radial fluorescence distributions

Radial profiles were calculated using radial_profile.py. For each client-positive ACAN condensate, the boundary was defined by the half-maximum ACAN intensity contour: background + 0.5 × (peak − background). Pixels were assigned a normalized radial coordinate ρ, with 0 at the center, 1 at the boundary, and values >1 outside. Intensity-weighted mean radial positions were computed for ACAN and the client. Peripheral displacement was calculated as 100 × (mean ρclient/mean ρACAN − 1), with positive values indicating a more peripheral client distribution.

### Statistical analysis

Analyses were performed using GraphPad Prism v10.0, R, and Python as specified above. Data were displayed as mean ± SEM unless otherwise indicated. The original figure analyses used Student’s t tests for two-group comparisons, one-way ANOVA with Šidák or Tukey multiple-comparison procedures for selected multigroup comparisons, Wilcoxon rank-sum tests for nucleus-level module scores, and Kolmogorov–Smirnov tests for selected distributions. A p value <0.05 was used as the nominal significance threshold. The number of animals, cells, images, nuclei, condensates, and independent experimental repeats is indicated separately in the figure legends where available. Nucleus-level sequencing comparisons were exploratory because there was one library per condition.

---

**Figure S1.**
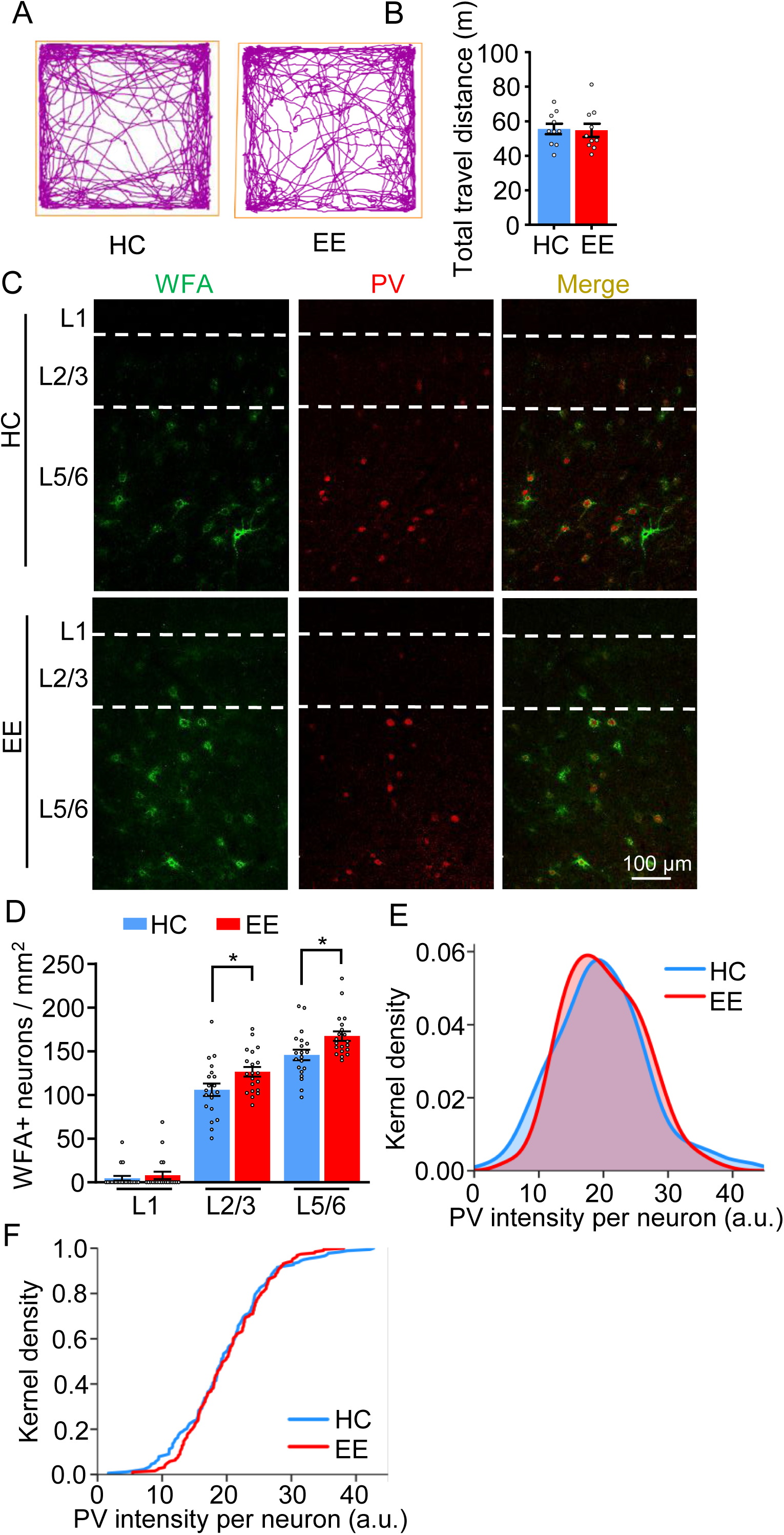
Locomotor Activity, WFA+ Neuron Density, and PV Intensity in HC and EE Mice, Related to **Figure 1**. **(A and B)** No significant difference in locomotor activity in HC and EE mice. A, representative travel traces in the open field test. B, total distance traveled. *n* = 11 mice per group. ns, not significant; unpaired Student’s *t* test. **(C and D)** Increased density of WFA+ neurons across PFC layers in EE mice. C, representative images of WFA labeling and PV immunostaining in the PFC of HC and EE mice. Scale bar, 100 μm. D, density of WFA+ neurons across PFC layers. *n* = 20 images from 3 mice per group. \**p* < 0.05, one-way ANOVA. **(E and F)** Kernel density estimation (E) and cumulative distribution (F) plots of PV intensity per neuron in HC (blue) and EE (red) mice. Kolmogorov–Smirnov test, D = 0.096, *p* = 0.341. *n* = 178 neurons from 3 mice in HC; *n* = 190 neurons from 3 mice in EE.

**Figure S2.**
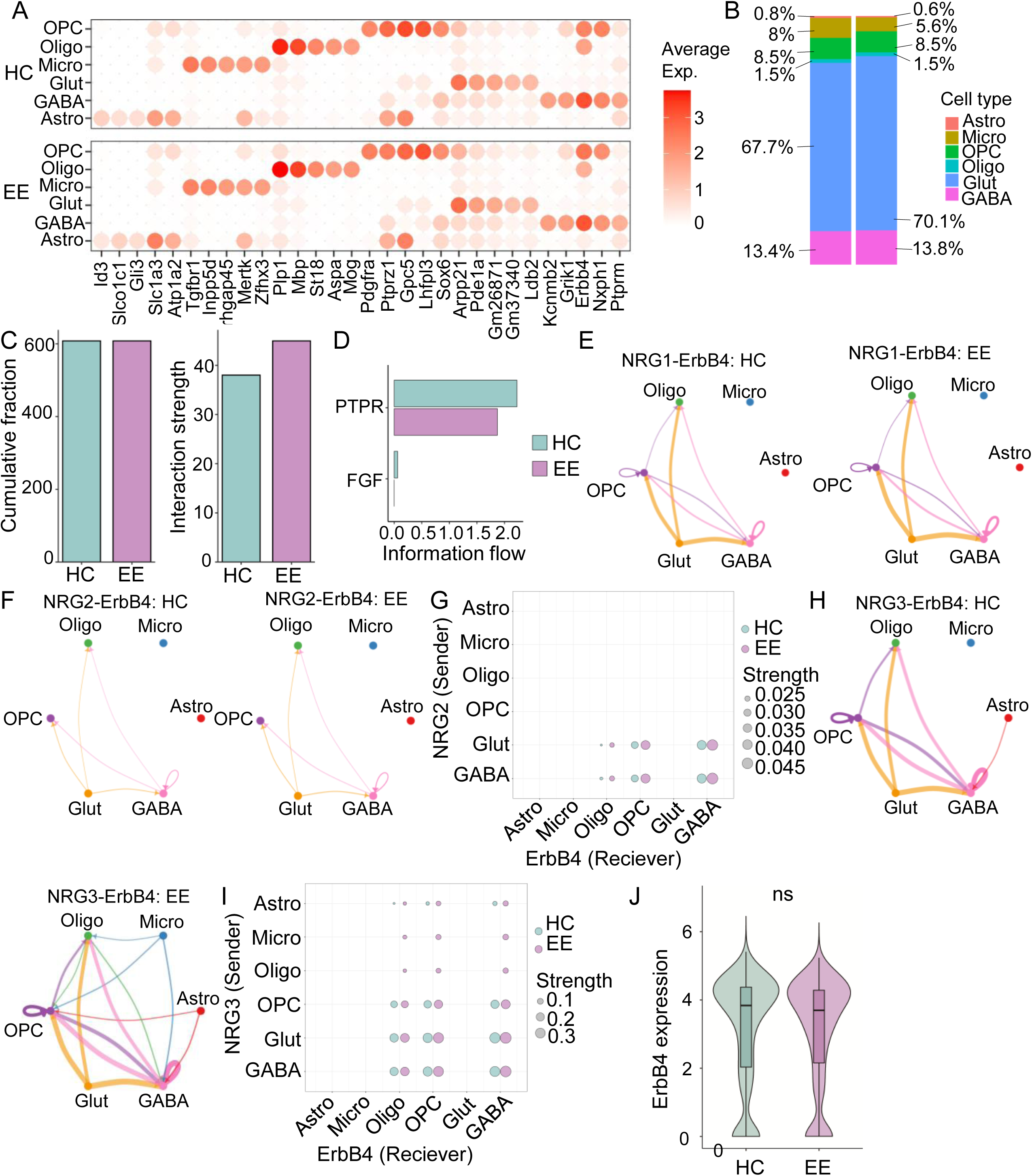
Cell-Type Annotation and Inferred NRG–ErbB4 Communication in HC and EE Groups, Related to **Figure 2**. **(A)** Marker-gene expression across annotated cell types. **(B)** Cell-type composition of HC and EE groups. **(C)** Number of inferred cell–cell interactions (left) and total inferred cell–cell communication strength (right) in HC and EE groups. **(D)** Inferred CCC pathways with higher communication probabilities in the HC group. **(E)** Cell-type-specific inferred NRG1–ErbB4 communication in HC and EE groups. **(F and G)** Inferred NRG2–ErbB4 communication in HC and EE groups. F, cell-type-specific inferred NRG2–ErbB4 communication. G, comparison of inferred NRG2–ErbB4 communication strength between groups. **(H and I)** Increased inferred NRG3–ErbB4 communication after EE. H, cell-type-specific inferred NRG3–ErbB4 communication. I, comparison of inferred NRG3–ErbB4 communication strength between groups. \**p* < 0.05, one-way ANOVA. **(J)** Violin plots of *Erbb4* transcript expression in HC and EE groups. ns, not significant.

**Figure S3.**
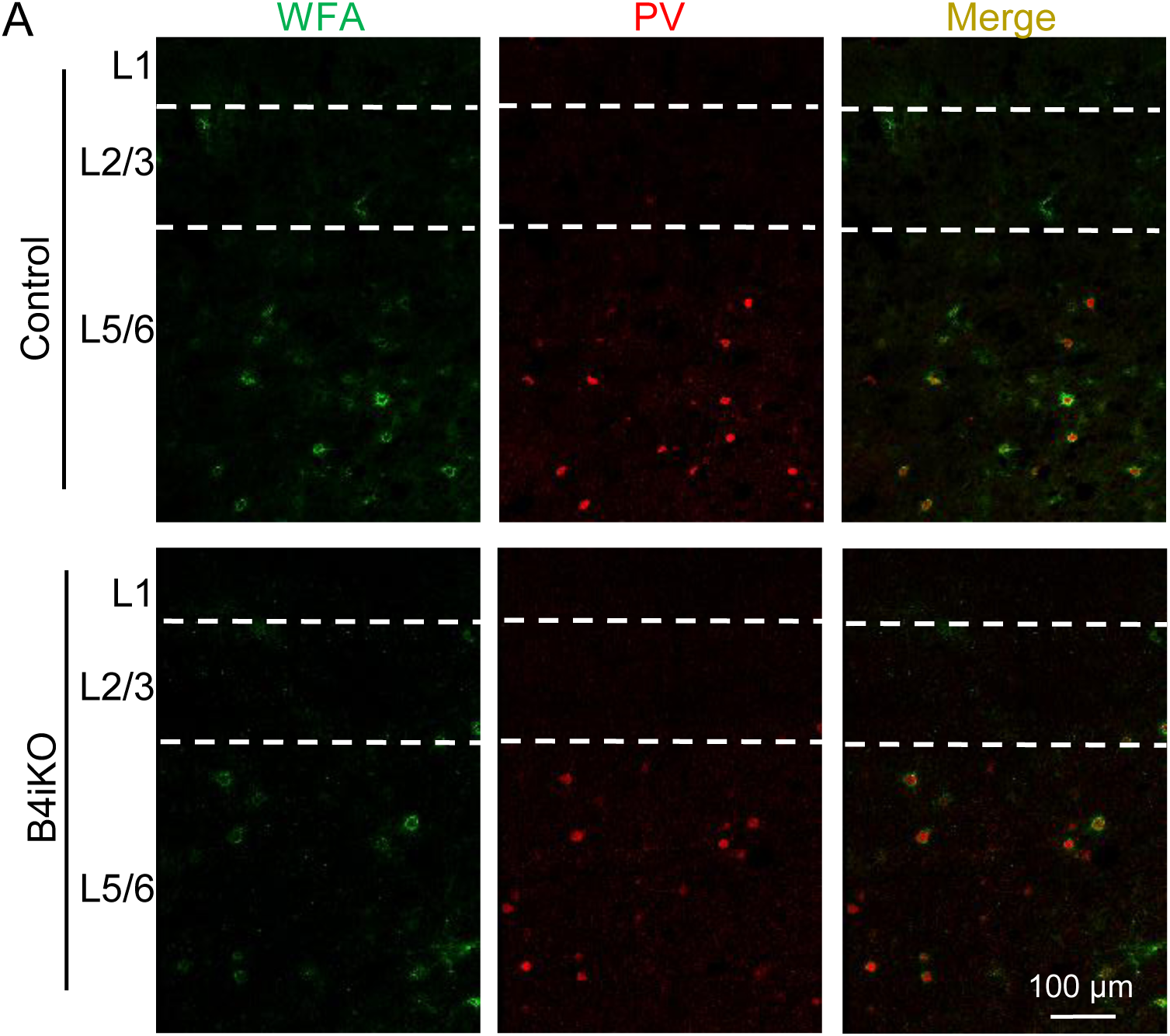
WFA and PV Labeling in the PFC after Inducible ErbB4 Deletion, Related to Figure 3. **(A)** Representative images of WFA labeling and PV immunostaining across PFC layers in control and B4iKO mice. Scale bar, 100 μm.

**Figure S4.**
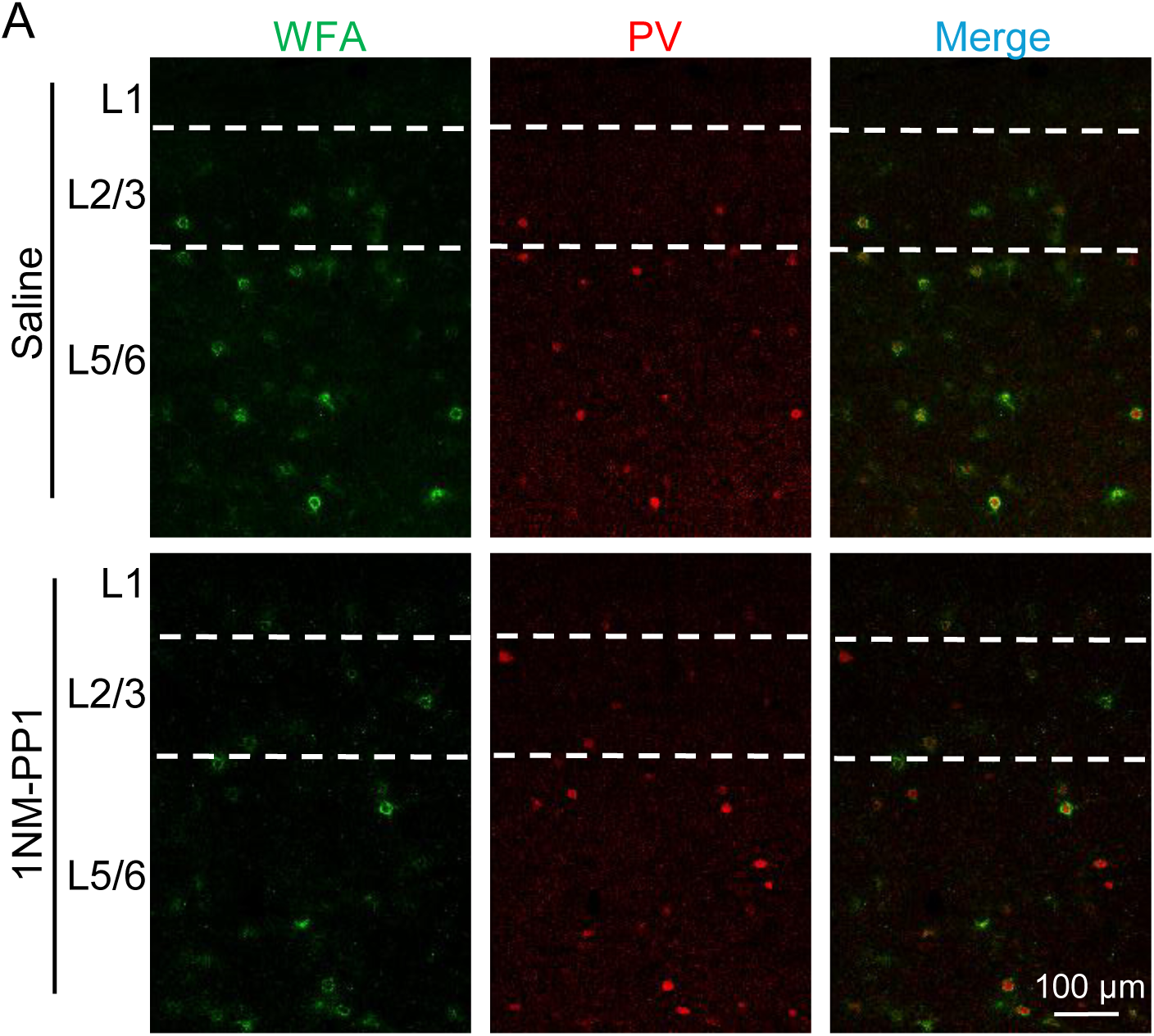
WFA and PV Labeling in the PFC of Saline- and 1NM-PP1-Treated ErbB4-T796G Mice, Related to Figure 4 **(A)** Representative images of WFA labeling and PV immunostaining across PFC layers in saline- and 1NM-PP1-treated ErbB4-T796G mice. Scale bar, 100 μm.

**Figure S5.**
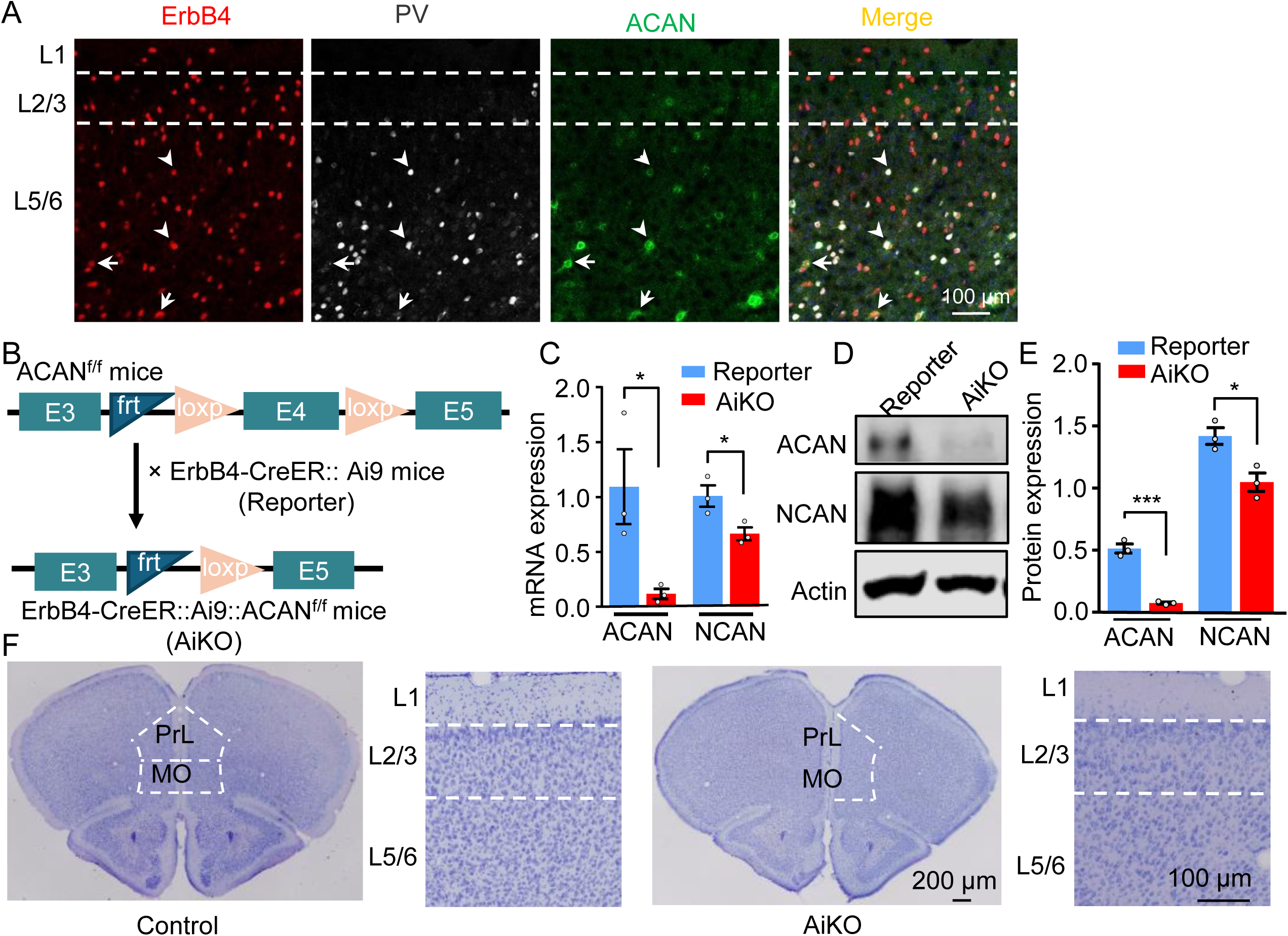
Validation of Conditional *Acan* Deletion and PFC Histology, Related to **Figure 6**. **(A)** Representative images of PV and ACAN immunostaining together with ErbB4 reporter fluorescence in the PFC of reporter mice. Arrows indicate ErbB4-reporter+/ACAN+/PV-neurons; arrowheads indicate ErbB4-reporter+/ACAN+/PV+ neurons. Scale bar, 100 μm. **(B)** Schematic of the *Acan* floxed allele and conditional recombination strategy used to generate ErbB4-CreER;Ai9;*Acan*^f/f^ (AiKO) mice. **(C)** Decreased mRNA levels of *Acan* and *Ncan* in AiKO mice. *n* = 3 mice per group. \**p* < 0.05, unpaired Student’s *t* test. **(D and E)** Decreased ACAN and NCAN protein expression in AiKO mice. D, representative immunoblots. E, quantification of the immunoblots in (D). *n* = 3 mice per group. \**p* < 0.05, \*\*\**p* < 0.001; unpaired Student’s *t* test. **(F)** Representative Nissl-stained PFC sections from reporter and AiKO mice. Scale bars, 200 μm (overview) and 100 μm (enlarged images). Data are presented as mean ± SEM.

**Figure S6.**
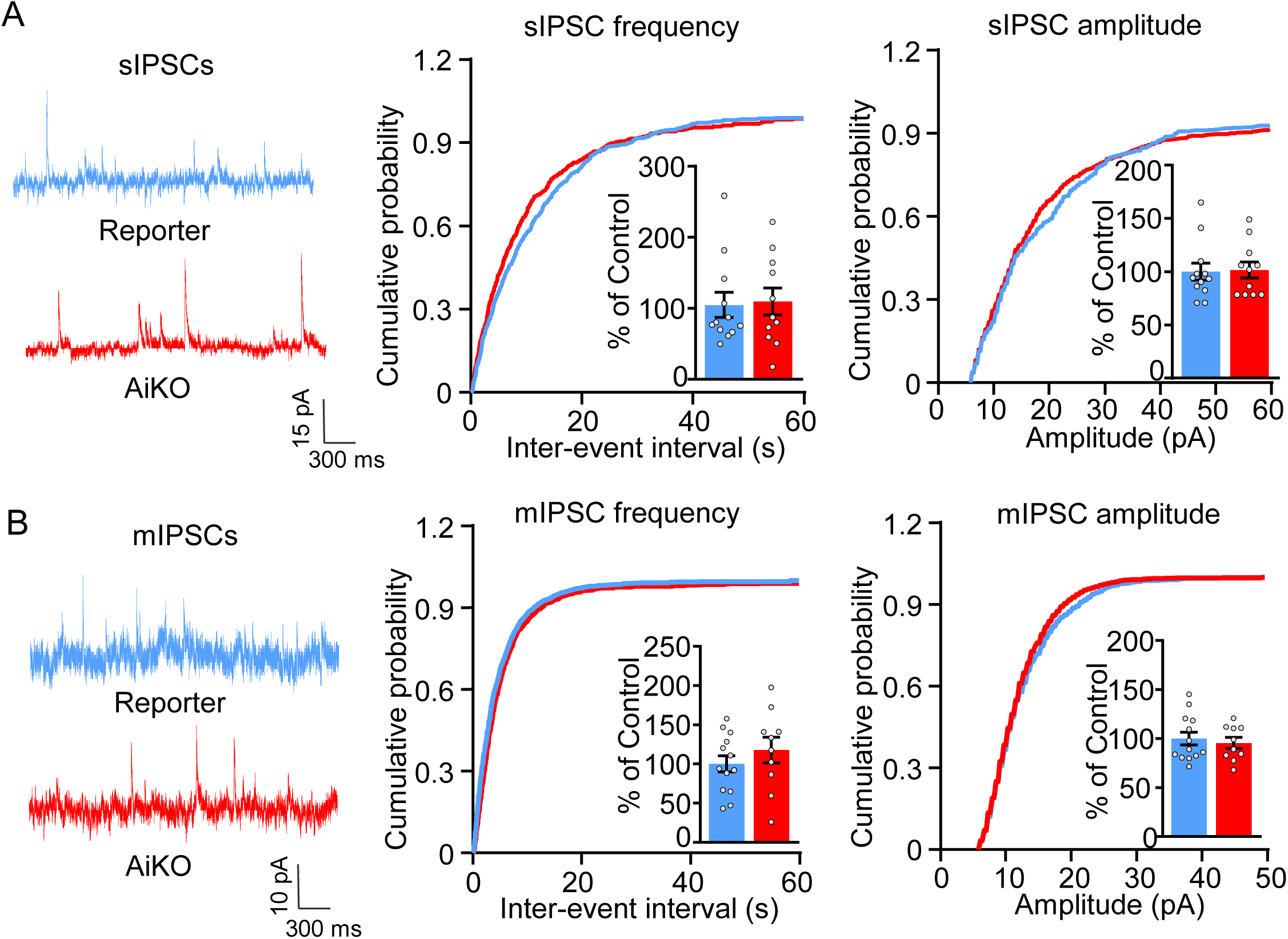
Inhibitory Synaptic Currents in Reporter and AiKO Mice, Related to **Figure 6**. **(A and B)** No significant difference in inhibitory synaptic transmission onto ErbB4+ interneurons in AiKO mice. A, sIPSCs; *n* = 12 neurons from 3 reporter mice and 11 neurons from 3 AiKO mice. B, mIPSCs; *n* = 13 neurons from 3 reporter mice and 10 neurons from 3 AiKO mice. For each, representative traces (left), CDFs of inter-event intervals and amplitudes (right), and normalized frequency and amplitude measurements (insets) in reporter and AiKO mice. Scale bars, 15 pA, 300 ms (A) and 10 pA, 300 ms (B). Not significant; unpaired Student’s *t* test. Data are presented as mean ± SEM.

**Figure S7.**
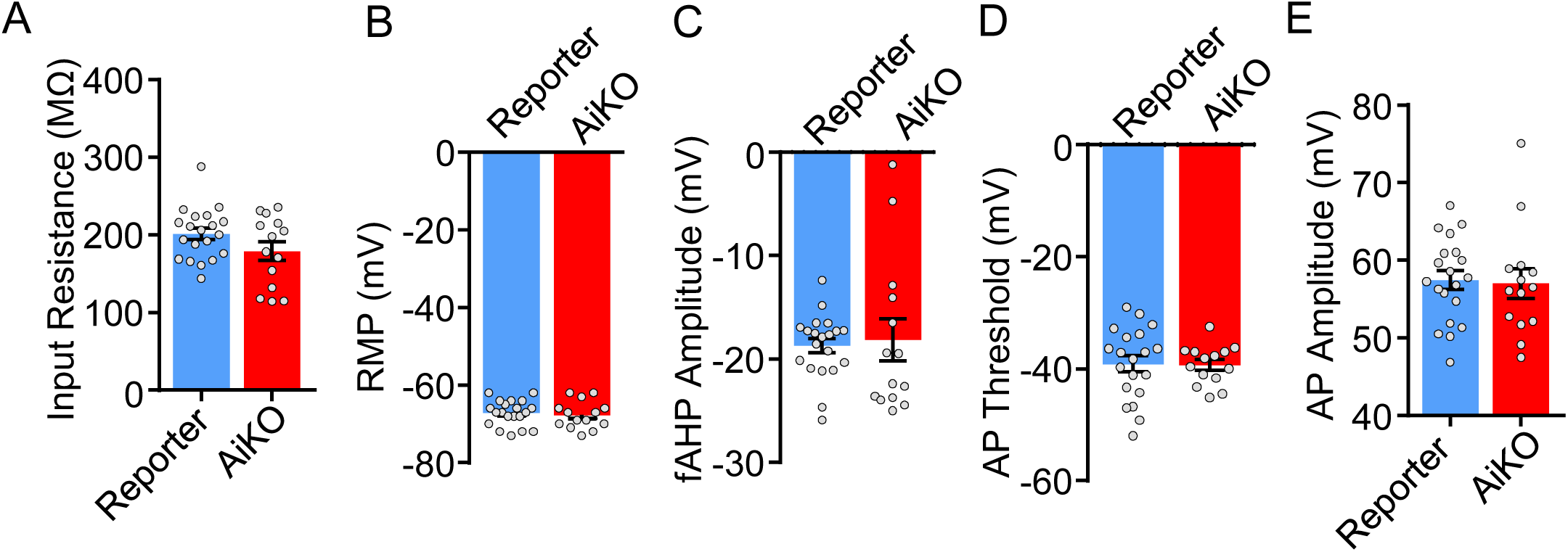
Intrinsic Membrane Properties of Interneurons in Reporter and AiKO Mice, Related to **Figure 6**. **(A-E)** No significant differences in intrinsic membrane properties in AiKO interneurons. A, input resistance. B, RMP. C, fAHP amplitude. D, AP threshold. E, AP amplitude. *n* = 14 neurons from 3 reporter mice; *n* = 20 neurons from 3 AiKO mice. Data are presented as mean ± SEM. Not significant; unpaired Student’s *t* test.

**Figure S8.**
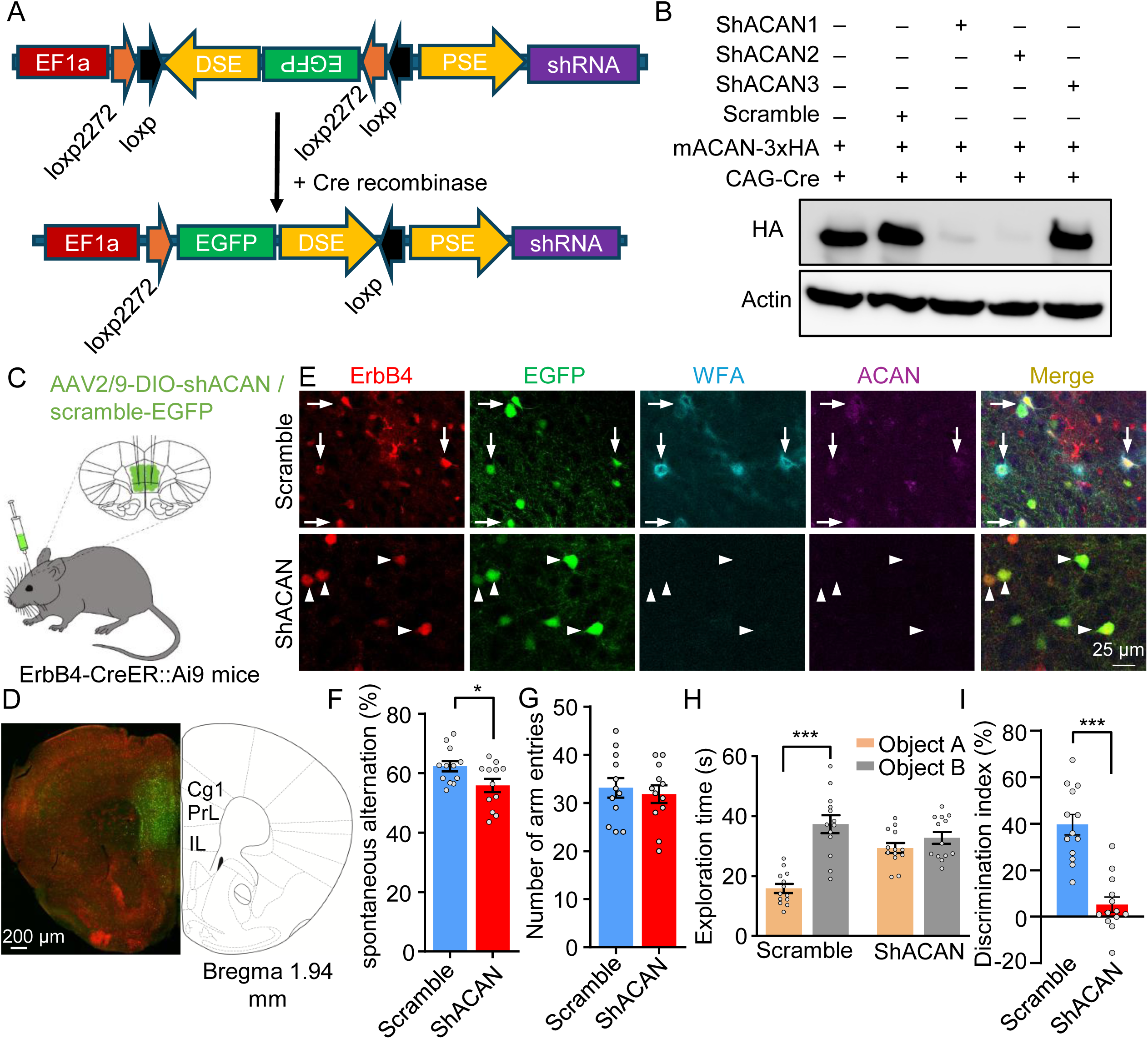
*Acan* Knockdown in ErbB4+ Neurons in the PFC and Memory Performance. **(A and B)** Design and validation of *Acan*-targeting shRNAs. A, schematic of the DIO-shACAN-EGFP constructs and the Cre-dependent strategy used to express shACAN and EGFP. B, representative immunoblot assessing knockdown efficiency of *Acan*-targeting shRNAs. **(C-E)** ACAN knockdown in ErbB4+ neurons impairs PNN expression. C, schematic of AAV stereotactic injection into the PFC. D, representative image of the injection site in the PFC of ErbB4-CreER;Ai9 mice (bregma 1.94 mm). Scale bar, 200 μm. E, representative images of ACAN immunostaining, WFA labeling, EGFP fluorescence, and the ErbB4 reporter signal in the PFC of AAV2/9-injected ErbB4-CreER;Ai9 mice. Scale bar, 25 μm. **(F and G)** Decreased spontaneous alternation in the Y-maze test in shACAN mice. F, spontaneous alternation in scramble and shACAN groups. G, no significant difference in the number of arm entries. *n* = 12 mice per group. \**p* < 0.05; unpaired Student’s *t* test. **(H and I)** Decreased discrimination index in the NOR test in shACAN mice. H, exploration time for objects A and B. I, discrimination index in scramble and shACAN groups. *n* = 13 mice per group. \*\*\**p* < 0.001; two-way ANOVA (H) and unpaired Student’s *t* test (I). Data are presented as mean ± SEM. See also Figures S9 and S10.

**Figure S9.**
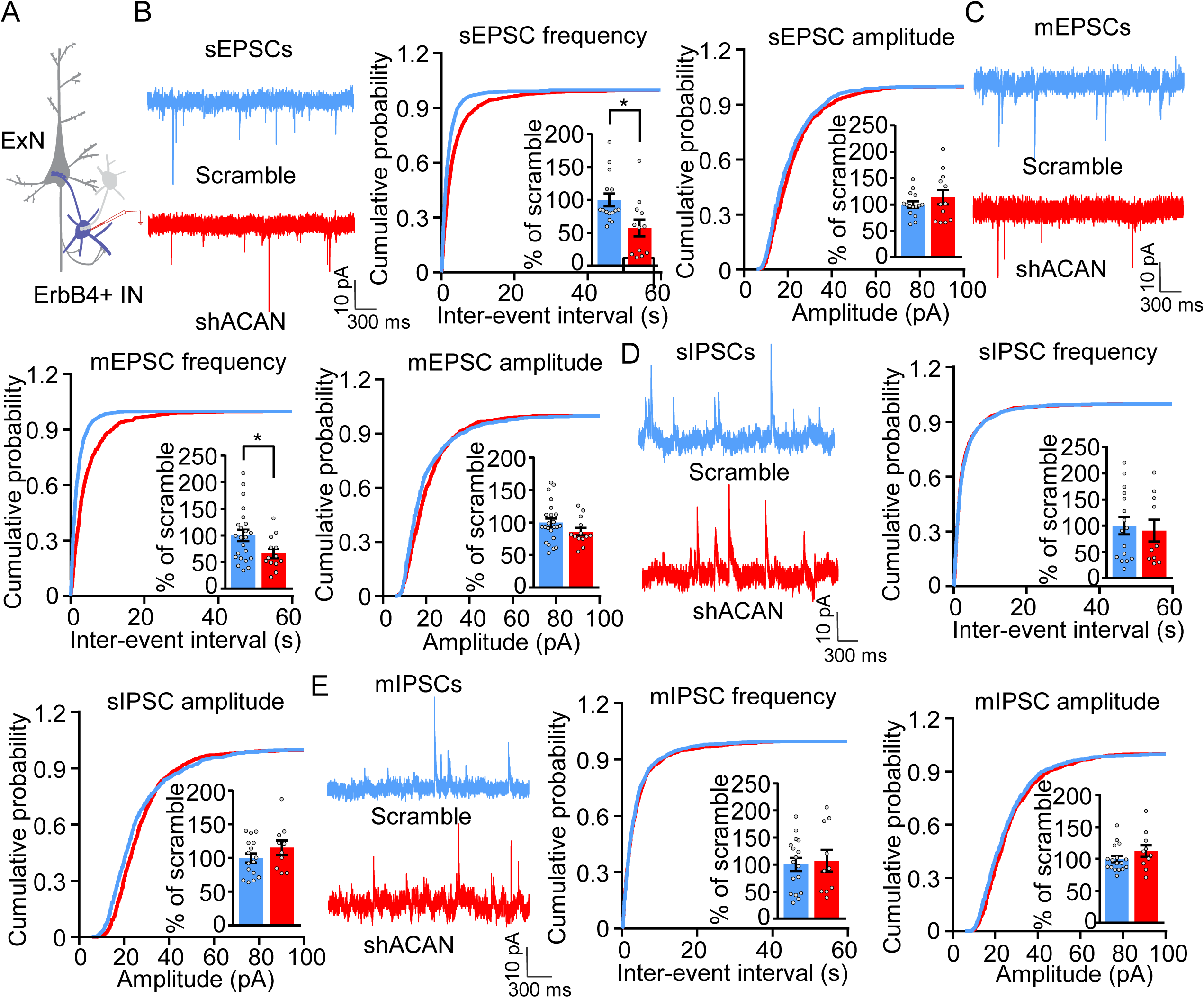
Synaptic Currents after *Acan* Knockdown in ErbB4+ Neurons in the PFC. **(A)** Schematic of whole-cell patch-clamp recording from ErbB4-targeted interneurons. **(B and C)** Decreased excitatory synaptic input onto ErbB4+ interneurons after *Acan* knockdown. B, the frequency and amplitude of sEPSCs; *n* = 15 neurons from 3 scramble mice and 12 neurons from 3 shACAN mice. C, the frequency and amplitude of mEPSCs; *n* = 23 neurons from 3 scramble mice and 13 neurons from 3 shACAN mice. For each, representative traces (left), CDFs of inter-event intervals and amplitudes (right), and normalized frequency and amplitude measurements (insets) in scramble and shACAN groups. Scale bars, 10 pA, 300 ms. \**p* < 0.05 for frequency, unpaired Student’s *t* test. **(D and E)** No significant difference in inhibitory synaptic transmission after *Acan* knockdown. D, the frequency and amplitude of sIPSCs; *n* = 16 neurons from 3 scramble mice and 10 neurons from 3 shACAN mice. E, the frequency and amplitude of mIPSCs; *n* = 17 neurons from 3 scramble mice and 10 neurons from 3 shACAN mice. For each, representative traces (left), CDFs of inter-event intervals and amplitudes (right), and normalized frequency and amplitude measurements (insets) in scramble and shACAN groups. Scale bars, 10 pA, 300 ms. Not significant, unpaired Student’s *t* test. Data are presented as mean ± SEM.

**Figure S10.**
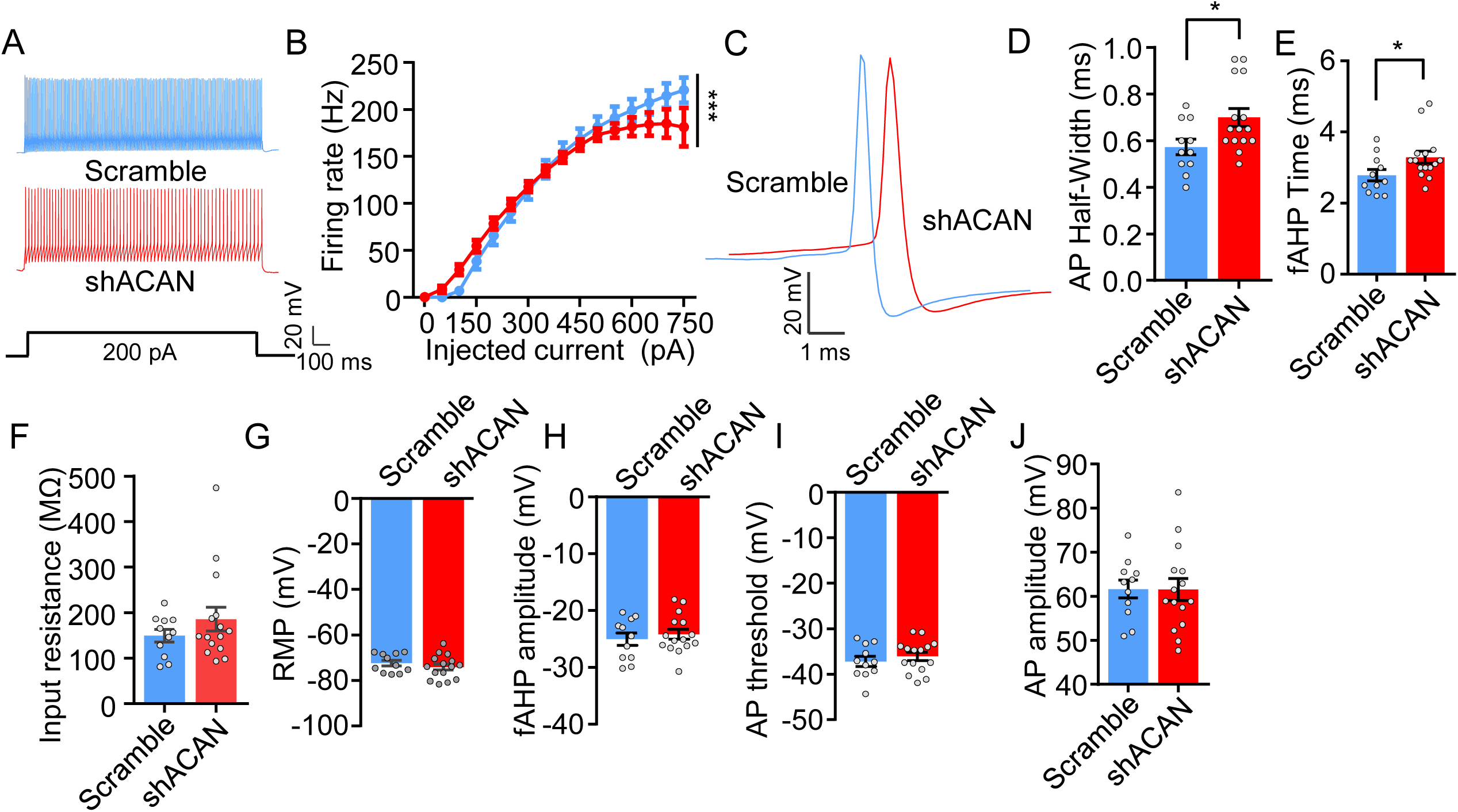
Intrinsic Membrane Properties after *Acan* Knockdown in ErbB4+ Neurons in the PFC. **(A and B)** Decreased firing rate of ErbB4+ interneurons in shACAN mice. A, representative traces of APs in scramble and shACAN mice. Scale bars, 20 mV, 100 ms. B, current injection–induced APs. *n* = 11 neurons from 3 scramble mice; *n* = 16 neurons from 3 shACAN mice. \*\*\**p* < 0.001, one-way ANOVA. **(C-E)** Broadened AP waveform and prolonged fAHP in shACAN mice. C, representative single APs of ErbB4+ interneurons from scramble and shACAN mice. Scale bars, 20 mV, 1 ms. D, AP half-width. E, fAHP duration. *n* = 11 neurons from 3 scramble mice; *n* = 15 neurons from 3 shACAN mice. \**p* < 0.05, unpaired Student’s *t* test. **(F-J)** No significant differences in intrinsic membrane properties in shACAN mice. F, input resistance. G, RMP. H, fAHP amplitude. I, AP threshold. J, AP amplitude. *n* = 11 neurons from 3 scramble mice; *n* = 15 neurons from 3 shACAN mice. Not significant; unpaired Student’s *t* test. Data are presented as mean ± SEM.

**Figure S11.**
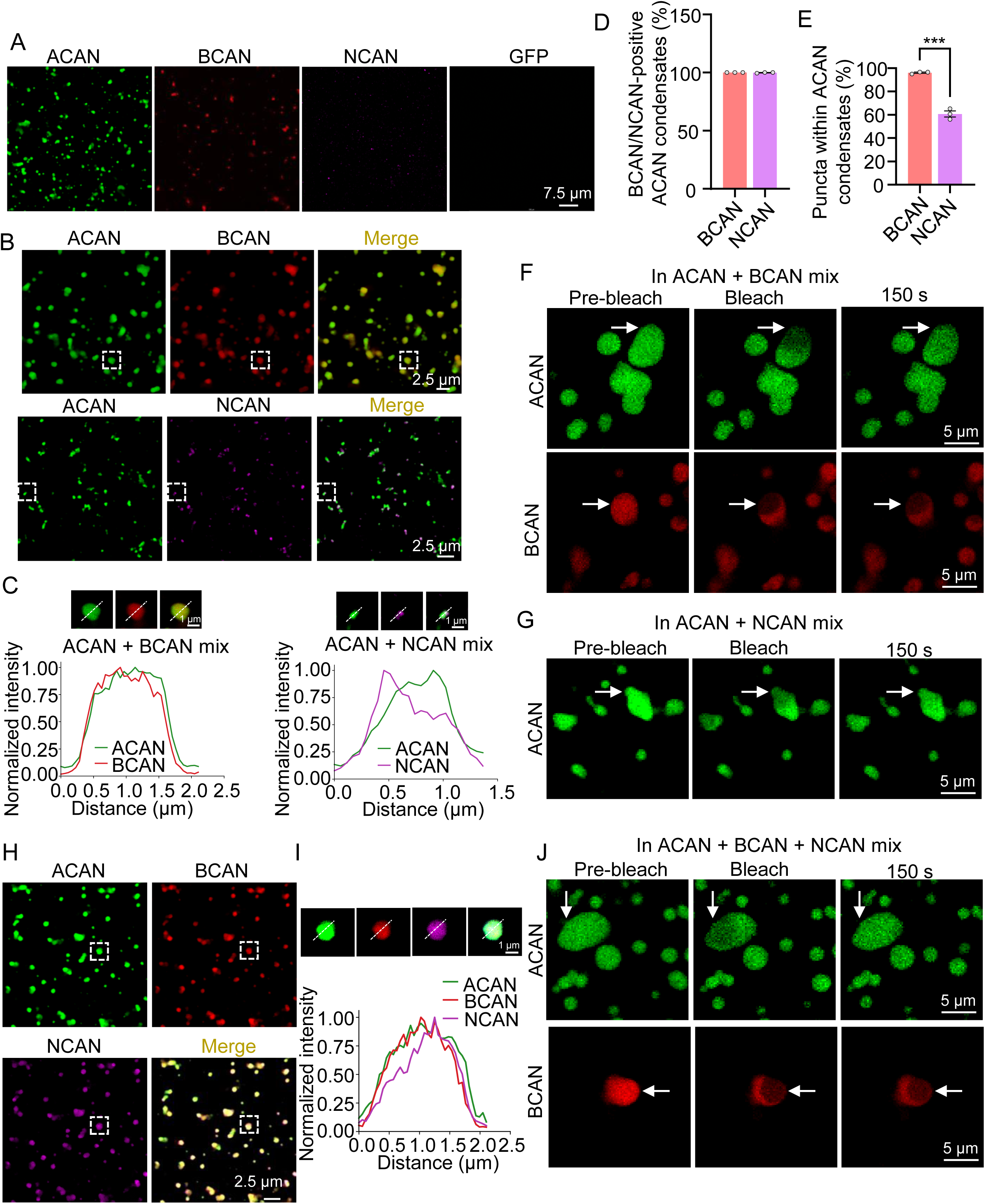
Co-condensation and FRAP of ACAN, BCAN, and NCAN, Related to **Figure 7**. **(A)** Representative overview fluorescence images of ACAN (green), BCAN (red), NCAN (magenta), and GFP, corresponding to Figure 7A. Scale bar, 7.5 μm. **(B)** Representative overview fluorescence images of ACAN + BCAN (upper panels) and ACAN + NCAN (lower panels) mixtures. ACAN, green; BCAN, red; NCAN, magenta. Scale bars, 2.5 μm. White dashed boxes mark the condensates enlarged in (C). **(C)** Fluorescence intensity profiles of ACAN (green) and BCAN (red) along a line crossing an ACAN-BCAN co-condensate or ACAN (green) and NCAN (magenta) along a line crossing an ACAN-NCAN co-condensate. Scale bars, 1 μm. **(D and E)** BCAN and NCAN are recruited into ACAN condensates. D, percentage of ACAN condensates positive for BCAN or NCAN, with the total number of ACAN condensates as the denominator (316 condensates from 7 fields for ACAN + BCAN; 658 condensates from 11 fields for ACAN + NCAN); each dot represents one independent preparation (mean of its fields of view); n = 3 preparations per group; not significant; unpaired Student’s *t* test. E, percentage of BCAN or NCAN puncta located within ACAN condensates, with the total number of BCAN or NCAN puncta as the denominator (271 BCAN puncta from 7 fields; 899 NCAN puncta from 11 fields); \*\*\**p* < 0.001; unpaired Student’s t test. **(F and G)** Representative FRAP images of ACAN + BCAN (F) and ACAN + NCAN (G) condensates before bleaching, immediately after bleaching, and at 150 s after bleaching, corresponding to Figure 7I and 7J. In (F), upper panels show ACAN and lower panels show BCAN. Arrows indicate the condensates subjected to photobleaching. Scale bars, 5 μm. **(H and I)** NCAN recruitment into ACAN condensates in the presence of BCAN. H, representative overview fluorescence images of ACAN + BCAN + NCAN mixtures. Scale bar, 2.5 μm. White dashed boxes mark the condensate enlarged in (I). I, fluorescence intensity profiles of ACAN (green), BCAN (red) and NCAN (magenta) along a line crossing an ACAN-BCAN-NCAN co-condensate. Scale bar, 1 μm. **(J)** Representative FRAP images of ACAN (upper panels) and BCAN (lower panels) in ACAN + BCAN + NCAN condensates before bleaching, immediately after bleaching, and at 150 s after bleaching, corresponding to Figure 7P. Arrows indicate the condensates subjected to photobleaching. Scale bars, 5 μm. Data are presented as mean ± SEM.

**Figure S12.**
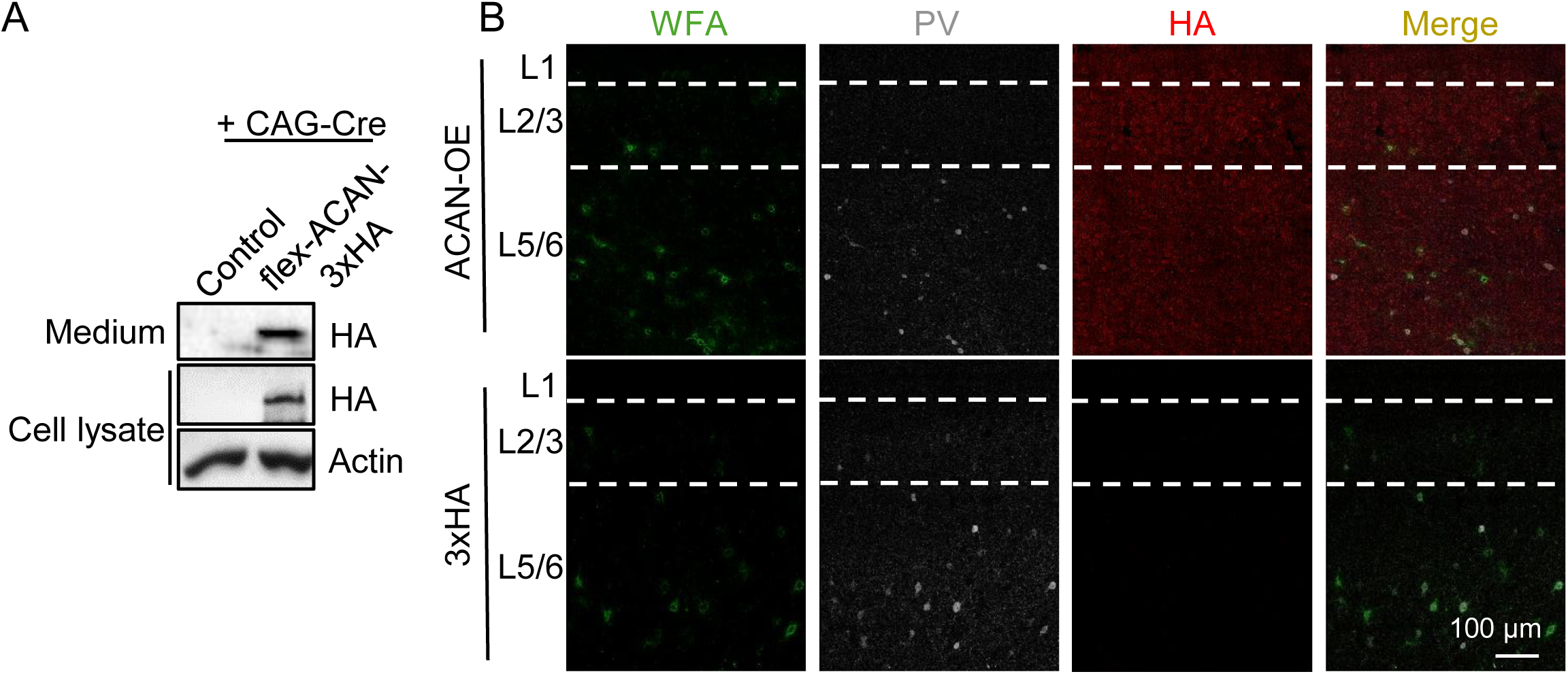
ACAN-3×HA Expression and WFA Labeling after Lentiviral Delivery, Related to **Figure 9**. **(A)** Representative immunoblots of ACAN-3×HA in transfected cell lysates and culture medium. **(B)** Representative images of WFA labeling and PV and HA immunostaining in the PFC of B4iKO mice following injection of the control or ACAN-OE lentivirus. Scale bar, 100 μm.

